# Disruption of TFIIH activities generates a stress gene expression response and reveals possible new targets against cancer

**DOI:** 10.1101/862508

**Authors:** Maritere Urioistegui-Arcos, Rodrigo Aguayo-Ortiz, María del Pilar Valencia-Morales, Erika Melchy-Pérez, Yvonne Rosenstein, Laura Domínguez, Mario Zurita

## Abstract

Disruption of the enzymatic activities of the transcription factor TFIIH by Triptolide (TPL) or THZ1 could be used against cancer. Here, we used an oncogenesis model to compare the effect of TFIIH inhibitors between transformed cells and their progenitors. We report that tumour cells exhibited highly increased sensitivity to TPL or THZ1 and that the combination of both had an additive effect. TPL affects the interaction between XPB and P52, causing a reduction in the levels of XPB, P52, and P8, but not other TFIIH subunits. RNA-Seq and RNAPII-ChIP-Seq experiments showed that although the levels of many transcripts were reduced, the levels of a significant number were increased after TPL treatment, with maintained or increased RNAPII promoter occupancy. A significant number of these genes encode for factors that have been related to tumour growth and metastasis. Some of these genes were also overexpressed in response to THZ1, which depletion enhances the toxicity of TPL and are possible new targets against cancer.

## Introduction

Cancer cells are known to be addicted to high levels of transcription as the enhanced expression of a plethora of molecules is required for the generation and the maintenance of a transformed phenotype (1, 2). This information suggests that the different factors that participate in general transcriptional activation by RNA polymerase II (RNAPII) could be targets for treating cancer. Since RNAPII is not able to recognize the promoter and initiate transcription in a regulated way by itself, it requires the assembly of what is known as the pre-initiation complex (PIC) at promoters. Generally, the PIC includes the TFIID complex, RNAPII, TFIIB, TFIIA, TFIIF, TFIIE, and TFIIH (3). In metazoans, during transcriptional activation, RNAPII synthesizes a transcript with a length of 20-120 nucleotides and then it pauses (4, 5). The release of paused RNAPII is conducted by the positive-elongation factor pTEFb (6, 7).

A component of the PIC and an interesting target to affect transcription— and therefore, cancer—is TFIIH (8, 9). TFIIH is a complex of 10 subunits composed of the CAK subcomplex containing CDK7, CYCH, and MAT1, which also participates in cell cycle control, and the core subcomplex that is part of the mechanism of nucleotide excision repair (NER) (10). The core subcomplex is composed of P8, P34, P44, P52, P60, XPB and XPD subunits (the last two are DNA helicases/ATPases) (11). Together, the CAK and the core form the 10-subunit TFIIH complex, which participates in transcription (11, 12). The role of TFIIH in RNAPII transcription involves phosphorylation of Ser 5 in the RNAPII CTD (p^Ser5^CTD RNAPII) by CDK7, which is important for transcription initiation, recruitment of the CAP enzyme, other modifications and mRNA processing (12–14). In contrast, XPB functions as an ATP-dependent translocase that rotates DNA to open it around the transcription initiation site, facilitating the synthesis of RNA by RNAPII (15, 16). Thus, compounds that affect the TFIIH functions have been found or developed as strong candidates to combat cancer. For instance, THZ1 and related compounds inhibit the kinase activity of CDK7 by binding a protein region outside of its catalytic domain (Cys312) (17) and is very effective in killing different types of cancer cells (18). On the other hand, triptolide (TPL), inhibits the ATPase activity of XPB by covalently binding its catalytic domain (19). TPL and its derivatives have been shown to kill different kinds of cancer cells (20). In addition, TPL has been used as a tool to study transcription initiation and promoter-proximal pausing (21–23). Although many studies demonstrate the potential use of TPL against cancer and indicate how this drug affects transcription initiation, studies of how TPL affects global gene expression between cancer cells and their progenitors are still needed. Additionally, it is not known whether TPL and THZ1 cause a similar effect in cells or whether TPL affects TFIIH integrity. In addition, transcriptional response studies are still limited or analysed with brief incubation times and not when the effect of TPL on cell homeostasis is initiating. In this work, we addressed these points using an inducible oncogenesis model.

## Methods

### Cell lines

MCF10A-ERSrc cell line was donated by Dr. Struhl. Cells were treated with 2µM tamoxifen (24) and morphological transformation is observed within 36-72 hours. MCF10A and MCF-10A-ER-Src cells were cultured in DMEM/F12 as previously described (24, 25) MDA-MB-231, MCF7 and HEK-293 cells lines were grown in DMEM (26, 27).

### Chemicals

Tamoxifen, 4-Hidroxytamoxifen (Sigma, Cat. H790); Triptolide (Tocris Cat. 3253) and THZ1 (APExBIO, Cat. A8882).

### Western blotting

Cell extracts were prepared as described in Gurrion et al, 2017 (28). Antibodies used were: 8WG16, H14, Phospho-STAT3, TBP, CDK7, MAT1, CYCLIN H, XPB, P52, P62, XPD, P8, EPAS, ID2, CRY2, HEXIM1, ACTIN and TUBULIN (Supplementary Table S1). Horseradish peroxidase (HRP)-coupled secondary antibodies (Invitrogen, 1:3000) were used for chemiluminescence detection through Thermo Scientific Pierce ECL.

### RT-PCR or qRT-PCR

Total RNA was extracted with TRizol (Invitrogen) and equal quantity from each sample were used. cDNA synthesis was performed in a reaction mix containing 1 µg of total RNA, oligo-dT, random primer and M-MLV Reverse Transcriptase (Invitrogen). For the intron cDNA same condition were used, except for 3 μg of total RNA and specific oligo. qPCR analyses were performed with LightCycler FastStart DNA Master^PLUS^ SYBR Green I and the LightCycler 1.5 Instrument (Roche). Relative expression level of each analysed gene was calculated by 2^-ΔCt^, where ΔCt= (Ct target gene - Ct control gene), using *GAPDH* as an internal control (29). Transcript abundance quiantification and DNA number copies were measured by triplicate and three independent biological replicates were analyzed. Primers are described in Supplementary Table S1.

### siRNA assays

siRNA-silencing was performed according to Dharmacon instructions. The siRNAs used were: *CRY2* (Cat. L-014151-01-0010), EPAS1 (Cat. L-004814-00-0010), *HEXIM1* (Cat. L-012225-01-0010), *ID2* (Cat. L-009864-00-0010), *LOC730101* (Cat. R-189565-00-0010) and Scrambled (Cat, D-001810-10-20). siRNAs were used in a 25 and 50nM final concentration for 24–72 hours and then mRNA or protein analysis were performed by western blot and flow cytometry.

### Flow cytometry

For proliferation assays, cells were loaded with the Cell Proliferation Dye eFluor 670 (Invitrogen, Cat. 65-0840). Viability and Apoptosis assays were according to manufacture: Fixable Viability Dye eFluor™ 780 (Invitrogen, Cat. 65-0865) and Biolegend Pacific Blue™ Annexin V or FITC Annexin V (BioLegend, Cat. 640918 or Cat. 640906, respectively). Intracellular protein staining was performed as previously described (30). Cell cycle assays were performed staining the cells with DAPI 5μg/ml in PBS for 30 min at 37°C. Samples were acquired on a BD FACSCanto II or BD FACs Aria Fusion flow cytometer with the BD FACSDiva software and analyzed using the FlowJo v10.5.3.

### Split-GFP assays

Plasmids pCNV_GFP1-9-OPT, pcDNA_GFP10-GCN4 and pcDNA_GCN4-GFP11, were donated by Dra. Cabantous (27). Stable transfection was performed in the HEK-293 cells of pCNV_GFP1-9-OPT plasmid using lipofectamine 3000. GCN4 sequences were removed with the restriction enzymes *Bspe*I:*Xba*I for pcDNA_GFP10-GCN4 and *Not*I:*Cla*I for pcDNA_GFP11-GCN4. *P52* (NM_001517) and *XPB* (NM_000122) were amplified from cDNA and inserted into *Mre*I:*Xba*I and *Not*I:*Cla*I cloning sites of pcDNA_GFP10 and pcDNA_GFP11 vectors respectively.

HEK-293_GFP1–9 cells were co-transfected with 1.5 mg of each plasmid: P52-GFP10 and GFP11-XPB. 36 hours after transfection, cells were visualized in the Olympus FV1000 Multi-photonic confocal microscope 60X. On the other hand, TPL at different concentration was added to co-transfected cells 8 hours later and 36 hours afterwards cells were stained for viability. As negative control the plasmids GCN4-GFP10 and GFP11-GCN4 with the same conditions were used.

### RNA-Seq and bioinformatic analysis

8X10^6^ cells were treated with TPL 125 nM or DMSO 4 h. Experiments were performed by duplicate. Total RNA was extracted with TRIzol (Invitrogen) according to manufacturer’s protocol. Poly-A enriched RNA was sequenced using Illumina HiSeq™ 2000 by the Beijing Genomics Institute (BGI). Briefly, mRNA enrichment was performed using oligo(dT), mRNA was fragmented and were used as template for the synthesis of cDNA by reverse transcription. The Agilent 2100 Bioanaylzer and ABI StepOnePlus Real-Time PCR System were used in quantification and qualification of the sample libraries. Libraries were sequenced in Illumina HiSeq™2000. Bowtie2 v0.9.6, (31) was used to map clean reads to reference gene and BWA v0.7.13 (32) to reference genome *hg19*. Sequencing reads were checked with FastQC v0.11.7. Expression levels were quantified using FPKM by RSEM v1.3.0 (33). Differential expression data was filtered using FDR ≤ 0.001, the absolute value of log_2_Ratio ≥ 1 and log FC ≥ 1.2. Gene expression analyses were performed using scripts in R v3.5.1. Analysis details are available upon request.

### ChIP-Seq and bioinformatic analysis

Cells were treated with TPL 125 nM or DMSO for 4 h. ChIP-Seq was performed according to a previously published protocol (34). Briefly, cells were crosslinked with 1% PFA at room temperature for 10 minutes and sonicated for 11 cycles (30 s ON/OFF, Diagenode Bioruptor Pico bath sonicator). 5% of the lysate was reserved as Input. Lysate was incubated with 5 mg of the antibody (8WG16) or irrelevant rabbit IgG (Invitrogen). Library construction and Illumina sequencing were performed using Illumina HiSeq SE50 platform at BGI.

Briefly, data filtering included removing adaptor sequences, contamination and low-quality reads from raw reads by BGI programs, like SOAPnuke filter., Clean data were mapped to the reference genome (*hg19*) by SOAPaligner/SOAP2 (35). MACS2 v1.4.2 (36) was used to call peaks and generate Bedgraph files that show fold change enrichment over input. Bedgraph files were then converted into BigWig files and uploaded to UCSC Genome Browser for visualization.

HOMER (Next-Generation Sequencing Analysis - *annotatePeaks.pl*) (37) was used to annotate the peaks to the genome. Difference in RNAPII enrichment between TPL-treated and control cells we performed a differential binding analysis using HOMER (getDiffExpression.pl). Differential expression data was filtered using log FC ≥ 1. Graphic representation of the data was performed using R v3.5.1 and GraphPad Prism v7. Analysis details are available upon request.

### Correlation of ChIP-Seq and RNA-Seq analysis

To perform a correlation between RNA-Seq and ChIP-Seq analyses we considerate only the RNAPII peaks located a ± 1 kb window spanning the Transcription Start Site (TSS). Differential expression data were filtered using log FC ≥ 1 for ChIP-Seq and log FC ≥ 1.2 for RNA-Seq. Graph was made GraphPad Prism (Fig 4C) Accession number for raw and processed ChIP-Seq and RNA-Seq data reported in this paper is GEO: GSE135256.

### Molecular dynamics

The cryo-electron microscopy structure of *Homo sapiens* TFIIH (PDB ID: 6NMI(38)) was retrieved from the Protein Data Bank (Fig 2F). The insolated XPB-P52-P8 submodule (17) as employed to perform the covalent docking of the optimized TPL structure into Cys342 residue of the XPB component employing the AutoDock v4.2 (39). The Apo and two holo (TPL-bound) forms of XPB-P52-P8 submodule were submitted to 100 ns all-atom molecular dynamics (MD) simulations using the AMBER99SB-ILDN force field (40) implemented in the GROMACS 5.1.4 package (41). (For more details, see Supplementary Methods and Supplementary Table S2.

## Results

### Triptolide and THZ1 preferentially kill transformed cells, and the combination has an additive effect

To study the effects of TPL and THZ1 on cancer cells and their progenitors, we used the MCF10A-ErSrc cell line as an oncogenesis model (24). After 36-72 h of incubation with tamoxifen, the MCF10A-ErSrc line is transformed, it is highly proliferative, has lost adherence, forms mammospheres, generates tumours and metastasis in immunocompromised mice. In this study, cells with these characteristics are referred to as <u>T</u>rans<u>F</u>ormed (TF) and their progenitors as <u>N</u>on-<u>T</u>rans<u>F</u>ormed (NTF). Phosphorylated STAT3 (pSTAT3) was used as a transformation control in this line (24, 42)

To analyse whether TF cells are more sensitive than NTF cells when TFIIH is affected, we evaluate by flow cytometry viability of NTF and TF cells which were incubated with TPL, THZ1 or both chemicals at different concentrations and for different times (Fig. 1A). The viability of both NTF and TF cells was highly affected by the presence of TPL, with TF cells being more sensitive (Fig. 1A). Following incubation with 100 nM TPL for 72 h, approximately 90% of TF cells died (Fig. 1A). However, at the same concentration and incubation time, more than 50% of NTF cells were still viable. In contrast, at a concentration of 250 nM THZ1 for 72 h, approximately 64% of TF cells died, and only 36% of NTF cells died (Fig. 1A). Interestingly, simultaneous incubation of NTF and TF cells with both chemicals had an additive effect on cell viability (Fig. 1A), as practically all TF cells died after 48 h when incubated with TPL (100 nM) and THZ1 (250 nM) (Fig. 1A). However, under those conditions and even after 72 h, approximately 40% of NTF cells remained viable (Fig. 1A). Thus, the combination of TPL and THZ1 is better than either drug alone and it preferentially targets TF cells for cell death. In all cases, cells died via apoptosis (Supplementary Fig. S1).

**Figure 1.**
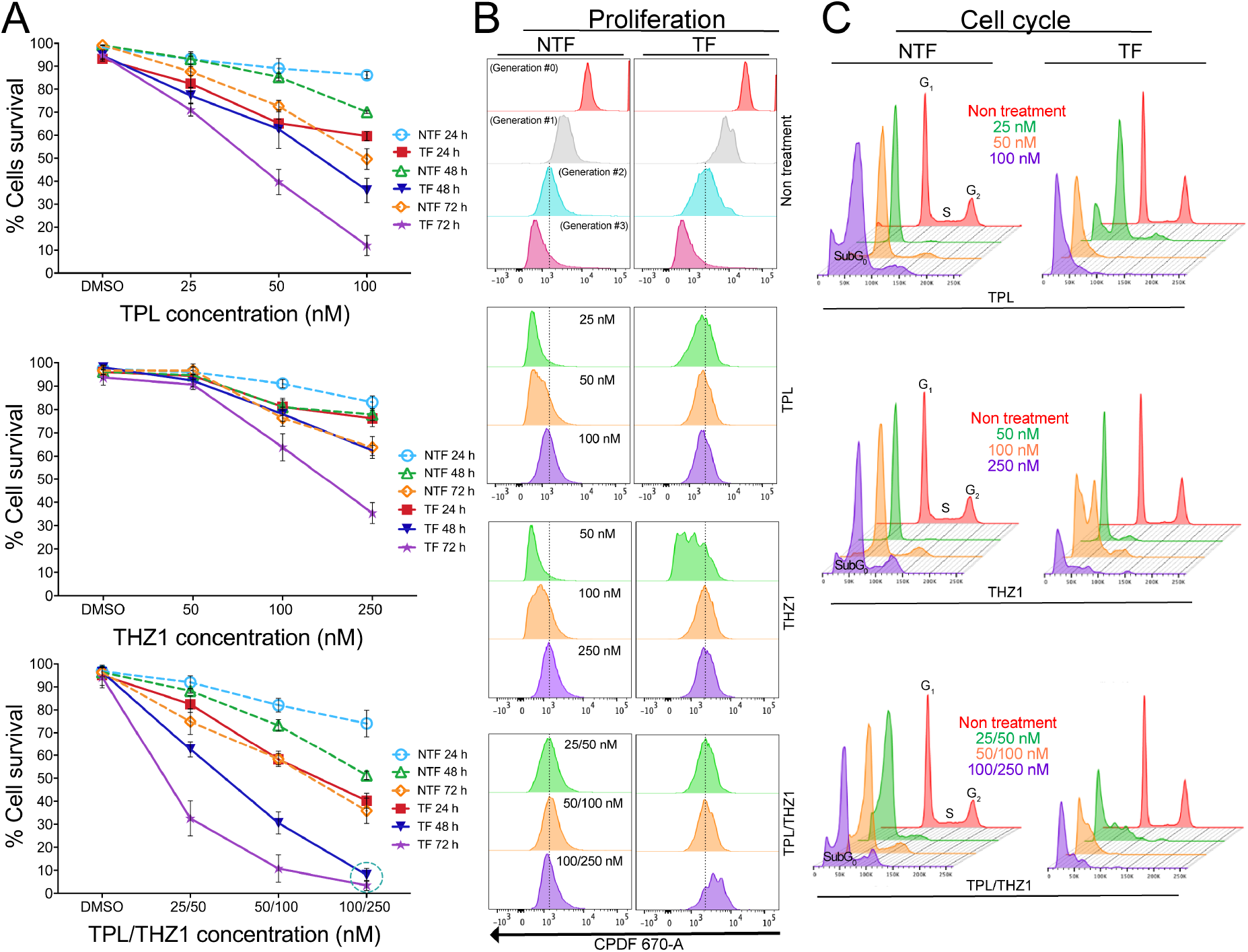
Triptolide (TPL) and THZ1 preferentially kill transformed cells and the combinatory of both substances potentiate its effect. **(A)** Cellular viability observed in transformed cells (TF) and non-transformed cells (NTF) incubated TPL (upper panel), THZ1 (midle panel) and the combination of THZ1 and TPL (lower panel). The concentrations used are indicated in the figure. **(B)** Proliferation assay in transformed cells (TF) and non-transformed cells (NTF) incubated TPL (upper panel), THZ1 (midle panel) and the combination of THZ1 and TPL (lower panel). The concentrations used are indicated in the figure. **(C)** Cell cycle arrest assay in transformed cells (TF) and non-transformed cells (NTF) incubated TPL (upper panel), THZ1 (midle panel) and the combination of THZ1 and TPL (lower panel). The concentrations used are indicated in the figure. In all the analysis the n=3.

Next, we evaluated by flow cytometry assays the effect of TPL, THZ1 and the combination of both at different concentrations and times on proliferation and cell cycle progression in NTF and TF cells (Fig. 1B-C). Figure 1B shows that after 72 h of incubation with 25 nM TPL, TF cells stopped after two cycles of proliferation; and that NTF cells required 100nM TPL to stop proliferating (Fig. 1B). Similarly, TF and NFT cells stopped proliferating, when incubated with 100 nM or 250 nM THZ1 for 72 h, respectively (Fig. 1B). Interestingly, when incubated with both TFIIH inhibitors NTF and TF cells stopped proliferating with only 25 nM TPL and 50 nM THZ1, confirming the additive effect of these drugs (Fig. 1B). Furthermore, we found that in the presence of TPL and THZ1, cells were arrested at the G_1_ phase and that lower concentrations of TPL and THZ1 were needed for TF cells (Fig. 1C). Taken together, these results indicate that TF cells are more sensitive to TPL and THZ1 than their NTF cells counterparts. TF cells stopped proliferating and were arrested at G_1_ at lower concentrations and shorter incubation times when incubated with either drug. Importantly, simultaneous treatment with TPL and THZ1 had a significantly more severe effect on TF cells than on NTF cells and than either drug used independently, underscoring the potential of simultaneously inhibiting different TFIIH activities with TPL and THZ to develop alternative therapies for cancer treatment.

### TPL interferes with the XPB-P52 interaction, inducing the degradation of the XPB-P52-P8 submodule of TFIIH

Based in our previous analysis of TFIIH mutants in *Drosophila* (43), we sought to explore the effect of TPL on the XPB levels in NTF and TF cells. Cells were incubated with 125 nM TPL for different times. Since the disruption of transcription by RNAPII causes degradation of this enzyme, we also evaluated levels of RNAPII as well as of other TFIIH subunits. As expected, levels of RNAPII—and therefore, p^ser5^CTD RNAPII—decreased as a result of incubating the cells with TPL (Fig. 2A). A clear reduction in XPB protein was also observed, which was greater in TF cells (Fig. 2A). Furthermore, levels of the P52 and P8 subunits were also diminished in response to TPL exposure (Fig. 2A). However, levels of other subunits of TFIIH, such as XPD, CDK7, CYCH, and MAT1, were not affected (Fig. 2A). As expected, incubation with THZ1 for different times reduced the levels of p^ser5^CTD RNAPII, but it did not affect the levels of this enzyme or the levels of any TFIIH subunit, including the CAK subcomplex, consistent with previous reports (44) (Supplementary Fig 2A).

**Figure 2.**
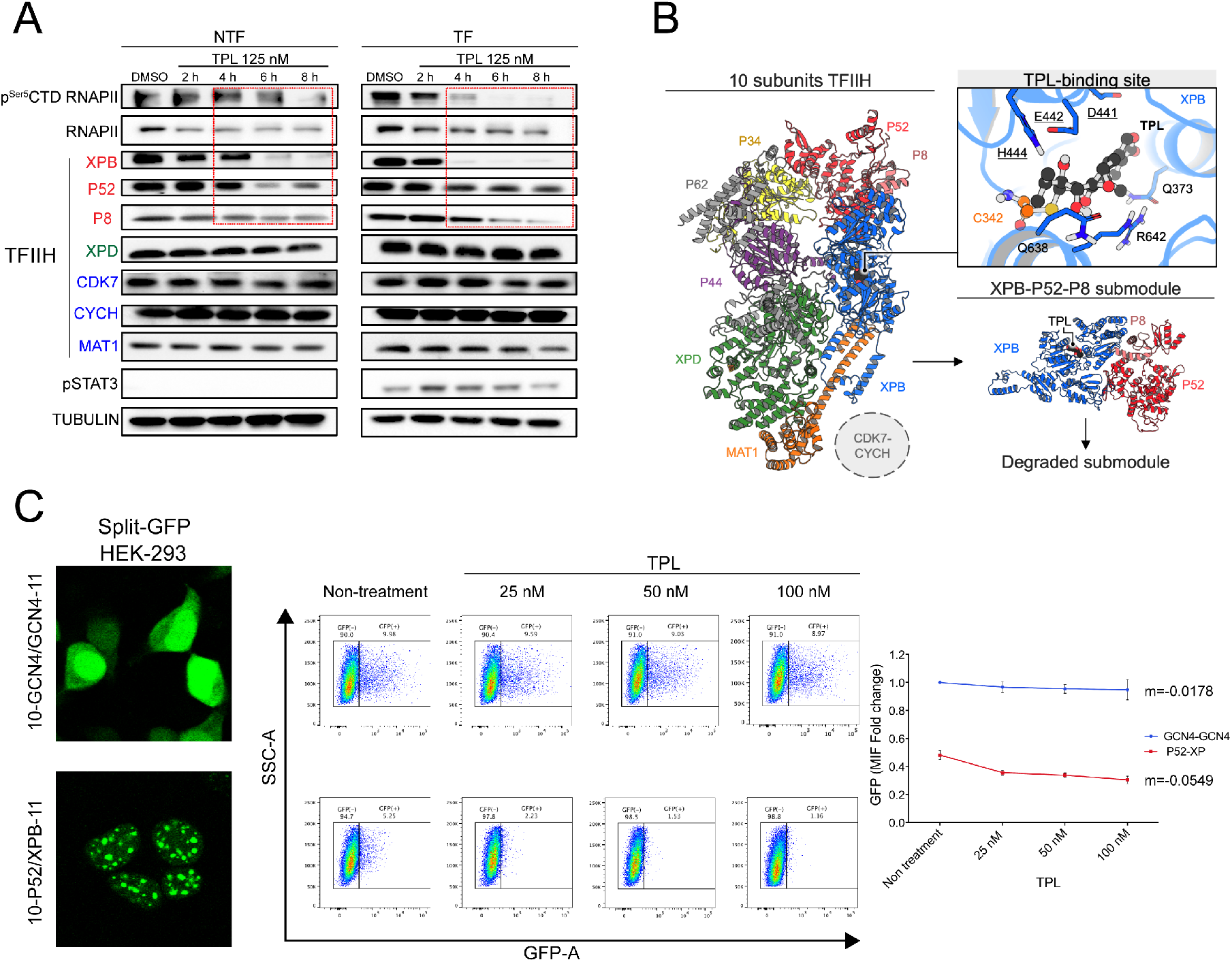
TPL induces the degradation of the XPB-P52-P8 submodule of TFIIH. **(A)** Western blots showing RNAPII CTD and the p^Ser5^ CTD RNAPII, and some TFIIH subunits (XPB, P52, P8, XPD, CDK7, CYCH, and MAT1) from cells incubated with TPL 125 nM for 2, 4, 6, 8 h in comparison with the control with DMSO for 8 h. Note that the TFIIH subunits, XPB, P52 and P8 protein levels decrease as the incubation time with TPL advances (box in red). The p-STAT3 is used as a transformation control in the line and tubulin protein as charge control. In all the analysis the n=3. **(B)** Computational molecular dynamics model that proposes the mechanism of the XPB-P52-P8 submodule dissociation from the TFIIH complex and degradation due to the covalent binding of TPL to XPB. The figure on the left shows the three-dimensional structure of TFIIH coloured by its components: XPB (blue), XPD (green), P8 (pink), P34 (yellow), P44 (purple), P52 (red), P62 (grey) MAT1 (orange). The upper right figure depicts TPL (black) binding mode in XPB. TPL binding site (TBS) residues are shown in blue, underlining the DEVH box amino acids, and C342 forming the covalent bond with TPL is highlighted in orange. **(C)** TPL interferes with the P52-XPB interaction. Split–GFP complementation assay between the P52 and XPB subunits expressed in HEK-293 cells are shown in the first panel. The second panel shows an example of the cytometry measure of the GFP fluorescent activity by the cell in the GCN4-GCN4 split GFP homodimer complementation used as control and the P52-GFP-XPB split complementation. Note that at 25 nM of TPL incubation by 28 h practically no fluorescent cells in the P52-XPB construct are detected. The right panel shows a kinetic assay of three independent experiments. In the GCN4-GCN4 control, the fluorescent remains constant with a negative pending of –0.01. However in the case of the P52-GFP-XPB fluorescence if has a negative pending of −0.05.

XPB directly contacts the P52 and P8 subunits and this interaction modulates the ATPase activity of XPB (43, 45). Since our results suggested that the binding of TPL to the ATPase domain of XPB destabilizes XPB as well as P52 and P8, we investigated whether TPL causes a distortion of XPB that limits the interaction of XPB with P52 and P8. To achieve this aim, we used the public information recently reported for the structure of the human TFIIH core by cryo-electron microscopy (38). TPL inhibits XPB ATPase function through the formation of a covalent bond between the C12 carbon (12,13-epoxide group) on the inhibitor and the sulfur atom of the Cys342 residue of XPB (TPL_C12_-Cys342) (46). The isolated XPB-P52-P8 putative submodule was employed to perform molecular dynamics (MD) simulations of the covalent docking of the optimized TPL structure to the Cys342 residue of XPB (Fig. 2B). Our covalent docking study showed that the TPL binding site (TBS) in XPB is located at the interface of the helicase domains (HD1 and HD2), which are mainly constituted by DEVH box residues (Supplementary Fig. 3). During the MD simulations, the presence of TPL at the HD1-HD2 interface altered the number of contacts between both domains, which may lead the separation of the domains and that the dissociation could be due to allosteric modulation guided by the loss of interactions between the XPB N-terminal domain (NTD) and P52 and between XPB HD2 and P52/P8.

To confirm the MD results we performed split-GFP-complementation assays between XPB and P52 by using the tripartite split-GFP association system (27). Stable HEK-293 cells expressing GFP1–9 were co-transfected with GFP10-P52 and XPB-GFP11 constructs an incubated with TPL at different low concentrations for 28 h (Fig. 2C). As a control, we used a GCN4 homodimeric interaction (GFP10-GCN4 and GCN4-GFP11) previous reported (27). The GFP fluorescence signal was quantified only in living cells by flow cytometry. After the TPL treatment a clear reduction in the fluorescence is observed in the P52-GFP-XPB-complementation cells, but not in the control cells (Fig. 2C). These *in vivo* results are in agree with structural modelling results that suggest that TPL interferes with the binging between XPB and P52. Altogether, results of this section suggest that XPB, P52, and P8 form a submodule in the core of TFIIH and that TPL besides to inhibit the XPB-ATPase activity, also cause the XPB-P52-P8 destabilization without affecting the rest of the TFIIH subunits.

### Analysis of the transcriptome of TPL-treated cells shows an unexpected gene expression response

While analysing the transcriptome of TFIIH mutants in *Drosophila*, we previously reported that not all genes responded equally and that the transcript levels of numerous genes increase in TFIIH mutant tissues (47, 48). Therefore, to explore whether TPL also generates a differential effect on gene expression in TF and NTF cells, we analysed the transcriptomes of these cells after incubation with 125 nM TPL for 4 hours, a concentration found to reduce equally RNAPII levels by half in NTF and TF cells, to affect mildly cell viability (Fig. 3A-B).

**Figure 3.**
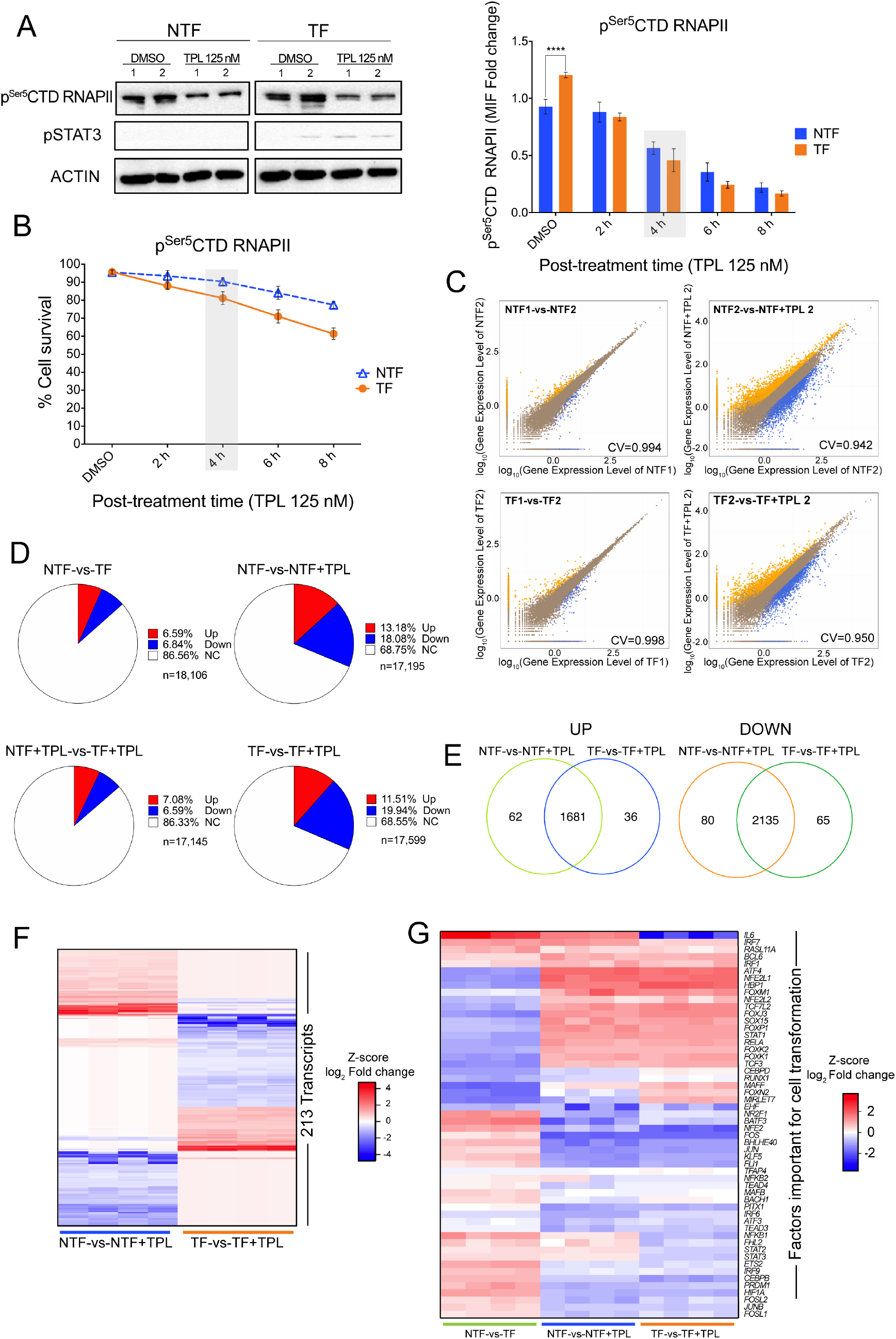
Transcriptome analysis of TPL treated cells. **(A)** Right panel show western blots to evaluate the p^Ser5^CTD RNAPII in non-transformed (NTF) and transformed (TF) cells treated with triptolide (TPL) for 4 h at 125 nM or DMSO for 4 h as control. pSTAT3 is used as transformation control and actin protein as control of loading (n=2). Left panel indicates the quantification of p^Ser5^CTD RNAPII by flow cytometry, NTF cells (blue) and TF cells (orange), the grey box, indicates the time that p^Ser5^CTD RNAPII has decreased approximately 50%. **(B)** Cell viability of the NTF cells (blue) and TF cells (orange) incubated with 125 nM of TPL for 2, 4, 6 and 8 h. DMSO (incubated for 8 h) were used as control. It can be appreciated that at 4 h (grey box) the cell viability is not very compromised (n=3). **(C)** Correlation between two of the replicas of the RNA-Seq data, (above and below left) and between two samples, one without treatment (DMSO control) and another incubated with TPL (4 h-125 nM); the x-axis and y-axis represent the log_10_ value of gene expression. The blue dots represent the down regulation transcripts; orange dots up-regulation transcripts and brown dots non-affected. **(D)** Percentages of up and down regulated genes in NTF, NTF cells treated with triptolide for 4 h at 125 nM (NTF+TPL), TF cells and TF cells treated with triptolide for 4 h at 125 nM (TF+TPL); in red are represented the transcripts percentage that increase, in blue the transcripts percentage that are down and in white transcripts that they with no significant log_2_ Fold change. **(E)** Venn diagram showing the percentage of up and down, unique and common transcripts between NTF and TF cells treated TPL. **(F)** Heat map comparing 213 transcripts that are differentially expressed between NTF and TF cells after TPL treatment. **(C)** Heat map showing the response to TPL in the expression of genes that are either up or down regulated in the establishment of the transformed phenotype. Note that TPL induce the increase in expression of most of the genes down regulated for the transformed phenotype and reduce the expression of must of the genes upregulated to maintain the transformed phenotype.

Approximately 18,500 different transcripts were identified in both TF and NTF cells (Supplementary Fig 4A). Correlation analysis between NTF and TF cells with and without TPL showed, as expected, that the treatment with TPL caused a reduction in the transcript levels of some genes, but intriguingly, the levels of other transcripts increased (Fig. 3C; Supplementary Fig 4B). Induction of the transformed phenotype of MCF10A-ErSrc cells reduced the expression of 6.84% and increased the expression of 6.59% of the genes (Fig. 3D). When we compared the effect of TPL in NTF and TF cells, the RNA levels of approximately 68% of the genes did not significantly change, probably because in the conditions used in this experiment, the reduction in transcription initiation of many genes is not detected by RNA-Seq (Fig. 3D). However, in both, NTF and TF cells, approximately 11% of the gene transcript levels were increased, and approximately 19% were decreased (Fig. 3D). Among the downregulated genes, 2135 transcripts were shared between NTF and TF cells. Among the upregulated genes, 1681 transcripts were common to NTF and TF cells, 62 were exclusive to NTF cells, and 36 were exclusive to TF cells (Fig. 3E; Supplementary Table S3). The expression of 213 genes were exclusive to either NTF or TF cells (Fig. 3F). Some genes as *SOX9*, *RETL1* and *IGFBP3*, were downregulated in NTF cells and upregulated in TF cells in response to TPL. However, we also detected genes whose transcripts were upregulated in NTF cells and downregulated in TF cells, such as *FAM111B* and *F2R* (Supplementary Table S3). Intriguingly, most of the recently identified factors required to maintain the oncogenic state in TF cells (49), change its expression back to NTF condition, suggesting a partial reversion of TF to NTF phenotype (Fig. 3G).

To confirm the RNA-Seq results, a set of 14 randomly selected genes was analysed by RT-PCR, and exhibited the same behaviour as shown by the RNA-Seq data (Supplementary Fig 5A). In addition, we explored whether a similar response to TPL occurs in other breast cancer cell lines (MCF-7 and MDA-MB-231) (Supplementary Fig 5A). Twelve out of the fourteen genes had a similar behaviour across all cells tested (Supplementary Fig 5A), indicating that breast cancer cell lines have a similar gene expression response to TPL.

All together our data indicate that XPB inhibition by TPL differentially modulates many genes in NTF and TF cells. Unexpectedly, the RNA-Seq results indicate that even when TPL affects RNAPII transcription, some genes are upregulated in response to this insult. This result suggests that there are genes for which, transcription may continue or even increase under conditions in which the levels of RNAPII are reduced as well as the XPB-P52-P8 submodule.

### Promoter occupancy and elongating RNAPII are maintained in genes upregulated in response to TPL

The increase in the RNA levels of numerous genes in response to TPL can be a result of different factors, including enhancement of transcription and/or an increase in accumulation of some RNAs by reducing their degradation. Reports using TPL to analyse pausing at promoters have been described (50, 51). However, the effect of TPL on RNA levels under conditions when the level of RNAPII is reduced was not determined in any of these studies. Therefore, we sought to analyse the genomic occupancy of RNAPII by chromatin immunoprecipitation followed by next-generation sequencing (ChIP-Seq) in NTF and TF cells incubated with or without TPL under similar conditions as those used in the RNA-Seq experiments.

Overall, the metagenome analysis showed that occupancy of RNAPII on the promoters was higher in TF cells than in NTF cells. As expected, we observed a reduction in the occupancy of RNAPII in NTF and TF cells when treated with TPL, with the main peak that corresponds to paused RNAPII in TPL-treated cells displaced to the 5’ end of the transcription start site (TSS) (Fig. 4A). This data shows that most RNAPII accumulates in the initiation state (PIC) and is in agreement with previous results on the effect of TPL on RNAPII occupancy on gene promoters (50, 51). Also consistent with the existence of highly stable paused promoters (50), since promoters corresponding to paused RNAPII were still detected (Fig. 4A). However, in our experiments, we identified stably paused promoters in cells with a substantial reduction in the levels of RNAPII and the XPB-P52-P8 submodule of TFIIH. Additionally, some promoters exhibited an increase in RNAPII occupancy in NTF (5.42%) and TF (3.9%) cells incubated with TPL (Fig. 4B; Supplementary Table S4),

**Figure 4.**
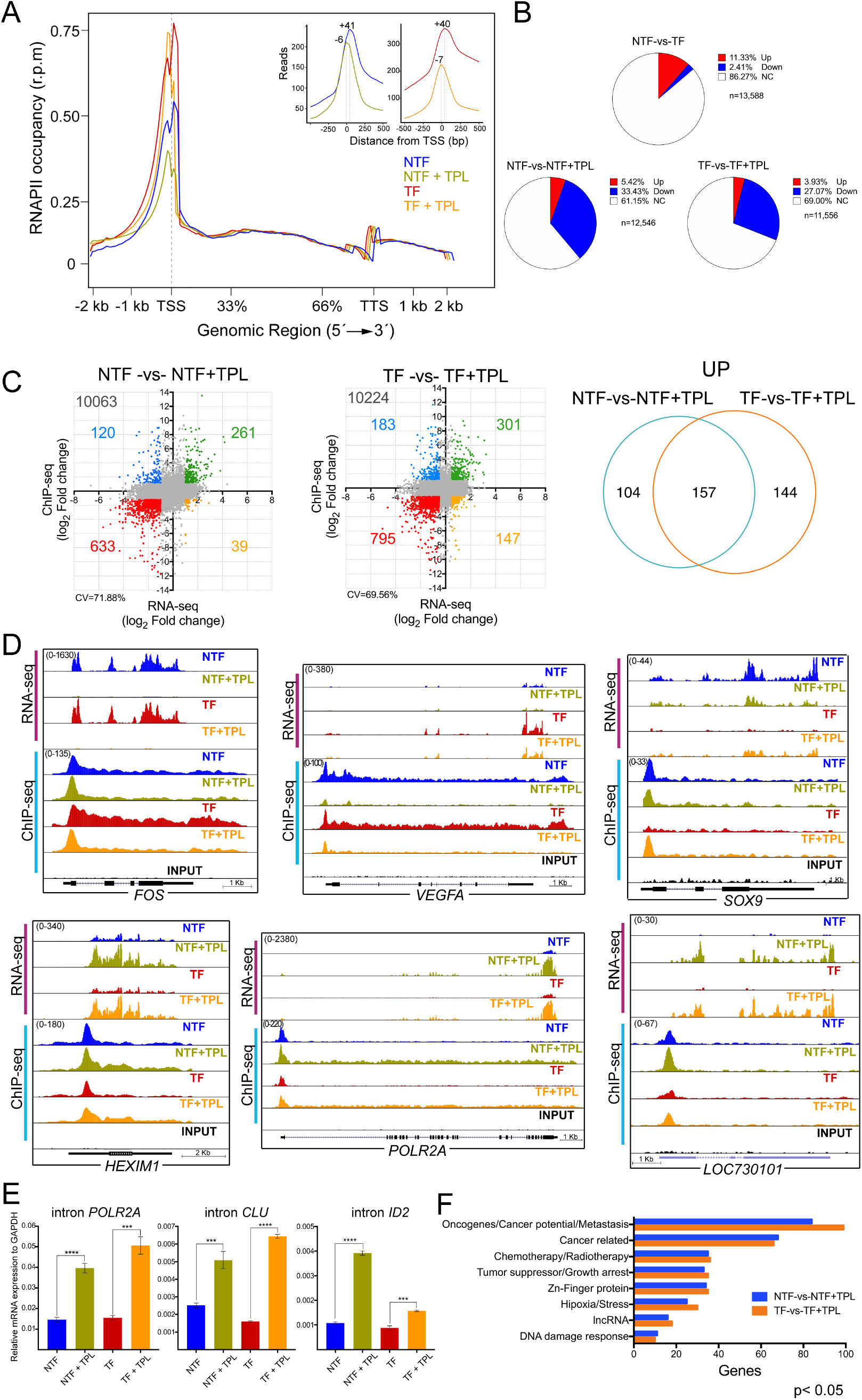
Analysis of the positioning of the RNAPII in cells treated with TPL. **(A)** Meta-analysis of the positioning of the RNAPII CTD comparing the non-transformed (NT, blue), non-transformed + triptolide (NTF, green), transformed (TF, red) and transformed + triptolide (TF+TPL, orange) treated with TPL for 4 hours at 125 nM. The upper right panel shows the displacement suffered by the cells when are treated with TPL towards the TSS-5’. Note the presence of two RNAPII picks, without TPL the major pick corresponds to paused RNAPII and with TPL the major pick corresponds to initiating RNAPII. Also note that after TPL treatment in some promoters RNAPII is maintained paused. **(B)** Relation of the occupancy of the RNAPII in gene promoters after the TPL treatment in NTF, NTF+TPL, TF and TF+TPL cells. In red is indicated the promoter occupancy percentage that increases, in blue percentage that goes down and in white promoters that do not have a significant change. **(C)** Correlation between the RNA-Seq data (x-axis) and ChIP-Seq data (y-axis), from the NTF-vs-NTF+TPL and TF-vs-TF+TPL cells. In green are genes that increase both in transcriptional level and in RNAPII positioning in the promoter; in blue genes that increase in transcriptional level but have low positioning in the promoter; in red are the double negative genes, genes with lower transcriptional level and lower positioning in the promoter and in yellow are genes with low transcript level but increase in the positioning of the polymerase. The right panel shows the number of genes that increase in RNA-Seq and in ChIP-Seq, unique and common between NTF-vs-NTF+TPL cells compared to TF-vs-TF + TPL. **(D)** Examples of the different behaviours observed in the RNA-Seq (pink bar) and ChIP-Seq (blue bar). Images are from the Genome Browser. **(E)** qRT-PCR of intronic sequences to quantify the levels de-novo synthetized mRNAs for genes over-expressed in response to TPL. POLR2A corresponds to the RNAPII large subunit gene, CLU is the clusterin gene and ID2 corresponds to the inhibitor of DNA binding gene. In the three genes an internal sequence of the first intron was evaluated. **(F)** Gene ontology of cancer related genes that were upregulated in the RNA-Seq and in the RNAPII occupancy in ChIP-Seq data.

Since the RNA-Seq results indicated that some transcripts were upregulated by TPL, we assessed whether there was a correlation between the occupancy level of RNAPII on the promoters with transcripts that were up- or downregulated in response to TPL. As shown in Figure. 4C, in TPL-treated NTF and TF cells, 261 and 301 genes respectively, were found to have an increased association of RNAPII to their promoters, and that correlated with a significant increase in their corresponding RNA levels (Supplementary Table S4). Of these genes, 157 were shared between NTF and TF cells (Fig. 4C). Also, an increase of RNAPII in the body of these genes is observed (Fig. 4D).

The correlation analyses indicated that there were different gene expression pattern in response to TPL (Fig. 4C). There were genes such as *FOS*, on which paused RNAPII was cleared, but initiating RNAPII was enriched and whose transcript levels were reduced by TPL in NTF and TF cells (Fig. 4D; Supplementary Table S4). For other genes, such as *SOX9*, the mRNA levels decreased in the presence of TPL in NTF cells, but in TF cells, TPL induced a significant increase in the RNA levels as well as an increase in RNAPII promoter occupancy of these same genes (Fig. 4D; Supplementary Table S4). The *VEGFA* gene was overexpressed in TF cells, with high levels of RNAPII in the body of the gene, but TPL repressed its expression, reducing the occupancy of RNAPII (Fig. 4D). The most intriguing response to TPL occurred in genes that were upregulated, such as *LOC730101* and *HEXIM1*, in which RNAPII occupancy on the promoter and along the body of the gene were increased (Fig. 4D). Interestingly, the effect of TPL on transcription induced potent overexpression of the gene that encodes for the large subunit of RNAPII (*POLR2A*) (Fig. 4D). We confirmed that the increase in levels of the mRNAs is due to an enhancement of transcription in response to TPL by evaluating the *novo* transcription of the first intron pre-mRNA of *POLR2A*, *CLU* and *ID2* genes by qRT-PCR (Fig. 4E).

Next, we performed a gene ontology analysis focusing on cancer related genes in which RNAPII occupancy on the TSS as well as its RNA levels was increased (Fig. 4F; Supplementary Table S5). A large number of genes that function as tumour and growth suppressors, such as *PHACTR4* and *ARID4A* (52, 53), as well as genes that correspond to chemotherapy/radiotherapy response genes, such as *SNAI1* and *SNX1* (54, 55), were overexpressed. However, a large number of genes considered as oncogenes or that promote tumour growth and/or metastasis, such as *SKI*, *EGR4*, *SNAI1* and *CEMIP2* (55–58), were also overexpressed in response to TPL (Fig 4F; Supplementary Table S4)

To confirm whether genes exhibiting an increase in RNAPII promoter occupancy were also overexpressed in response to TPL and for further analysis, we selected five genes with different cellular functions for qRT-PCR. The selected genes were *ID2*, a transcription factor known to participate in epithelial-mesenchymal transition (59); *CRY2*, a circadian repressor involved in MYC turnover (60); long noncoding RNA (lncRNA) *LOC730101*, which is induced by hypoxia and has been related to metastasis (61); and *HEXIM1*, an inhibitor of pTEFb, linked to chemotherapy resistance and that it has been shown that also is overexpressed in response to JQ1 (62, 63). In addition, we analysed *EPAS1*, also known as *HIF-2A*, a transcription factor that responds to hypoxia, a hallmark of cancer. *EPAS1* is overexpressed in TF cells and its transcript levels were high and maintained in cells incubated with TPL (59). For all genes, the increase in RNA levels was confirmed (Fig. 5A-E). A kinetic analysis of the encoded products of these genes by western blot confirmed that not only the accumulation of the RNA increase, but also the corresponding protein in response to TPL (Fig. 5F). Furthermore, we verified the increase in the expression of these genes in response to TPL in other breast cancer cell lines by RT-PCR. Interestingly, these genes were also overexpressed in the MCF-7 and MDA-MB-231 breast cancer cell lines in response to TPL, indicating that this response is not exclusive to the MCF10A-ErSrc line (Supplementarry Fig. 6). Collectively, our results indicate that both NTF and TF cells respond to a TPL insult by inhibiting RNA transcription. Yet, despite TPL directly affects transcription initiation, and significantly reduces the levels of RNAPII as well as of the XPB-P52-P8 submodule components of TFIIH, the transcriptional stress imposed to the cells results in activation of selected genes. Among those, genes that suppress cancer growth are overexpressed, but importantly, genes that promote carcinogenesis, chemotherapy resistance and metastasis are also upregulated.

**Figure 5.**
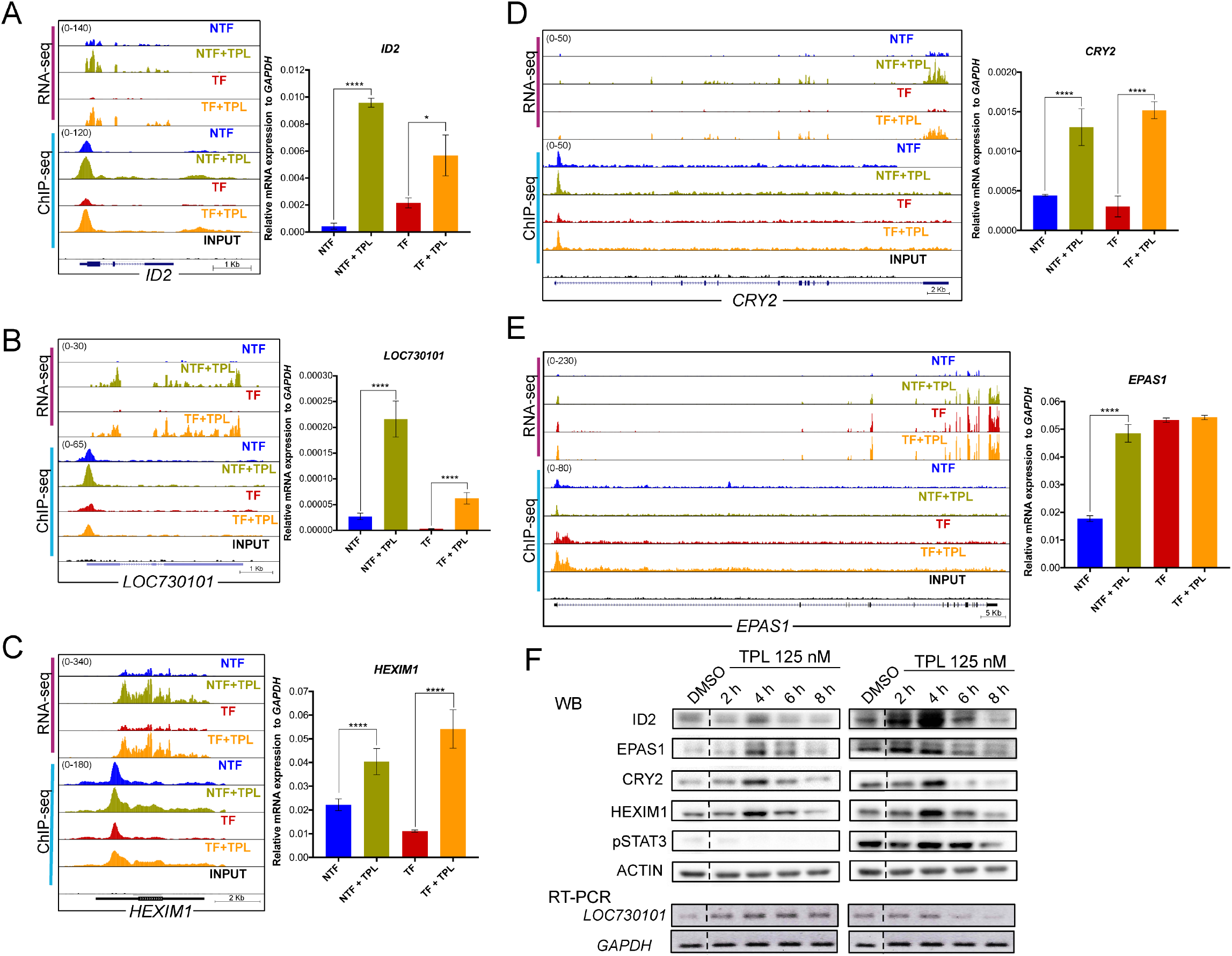
Analysis of overexpressed genes and its protein products in response to TPL treatment. **(A)** *ID2* gene, representation of the Genome Browser, RNA-Seq (pink bar) and ChIP-Seq (blue bar), comparing the NTF (blue), NTF+TPL (green), TF (red), TF+TPL (orange) and the input (black). qRT-PCR corroboration is also show. **(B)** lncRNA *LOC730101* representation in the Genome Browser. qRT-PCR experiments are also show confirming the increase of this transcript after TPL treatment. **(C)** *HEXIM* Genome Browser image. qRT-PCR experiments confirming HEXIM1 overexpression by TPL. **(D)** *CRY2* gene increase with the TPL treatment shown in the Genome Browser image, confirmation by qRT-PCR is also show. **(E)** *EPAS1* Genome Browser image and qRT-PCR corroboration **(F).** Time curse experiment of the overexpression of the ID2, HEXIM1, CRY2 and EPAS1 proteins in response to TPL by western blot analysis. A time course analysis of the overexpression of the LOC730101 lncRNA by RT-PCR is also show. STAT3 phosphorylated is an indicator of the transformed phenotype in TF cells

**Figure 6.**
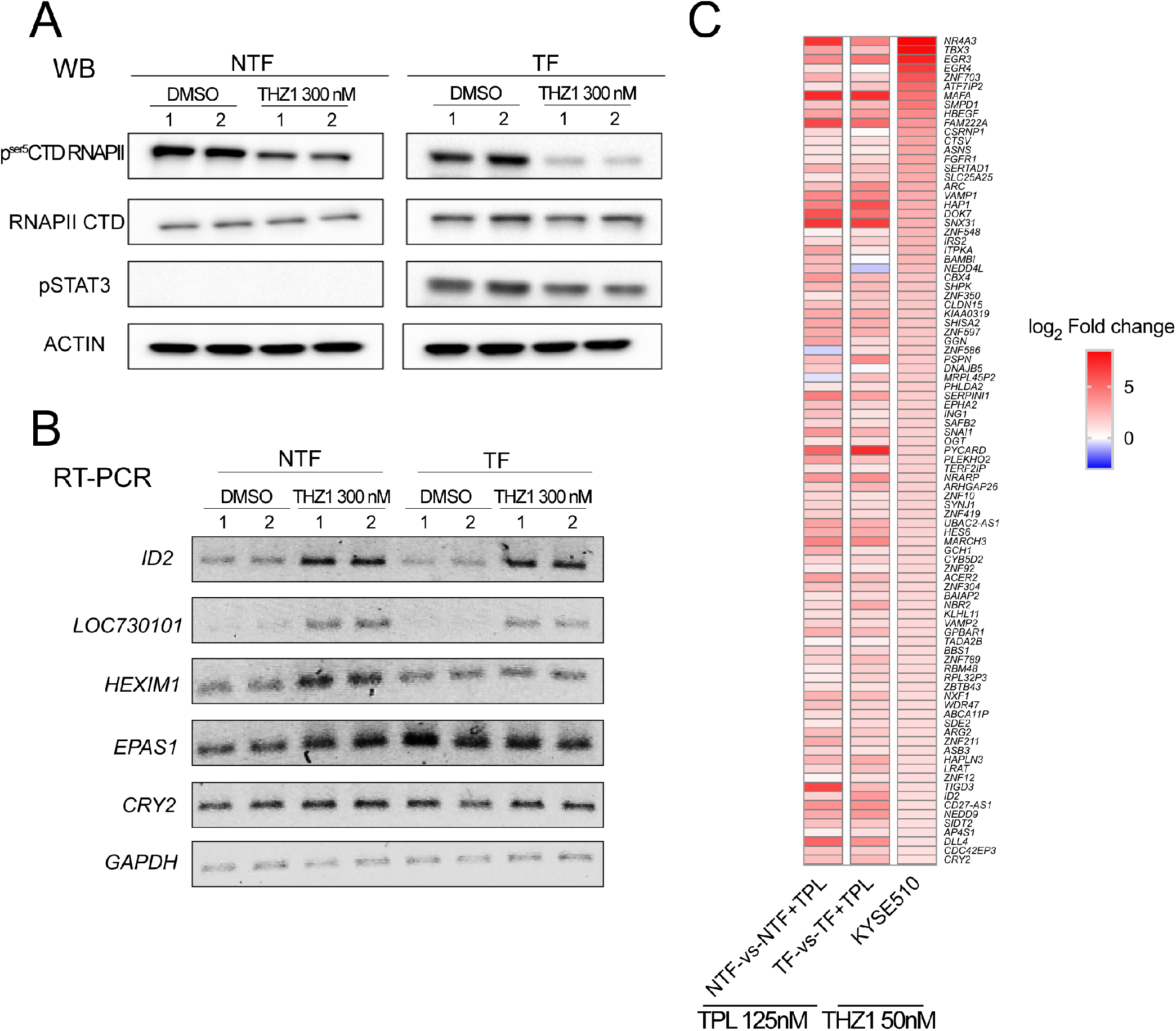
THZ1 generates a similar response as observed by the effect of TPL on transformed cells. **(A)** p^Ser5^CTD RNAPII and RNAPII CTD levels in NTF and TF cells treated with 300 nM of TPL for 2 h or with DMSO control as control. **(B)** Semiquantitative RT-PCR of NTF and TF cells treated with 300 nM of THZ1 for 2 h of *ID2, LOC730101, CRY2, EPAS1* and *HEXIM1*. *GAPDH* was used as load control. **(C)** Comparison with published data of the THZ1 response in KYSE510 cells (64) indicating a similar response to some of the overexpressed genes after TPL treatment.

### THZ1 drives a similar gene response as TPL in cancer cells

Next, we decided to explore whether incubation with THZ1 induces a similar gene response as TPL. The first approach to answer this question was to determine, by RT-PCR, the response of some of upregulated genes in response to TPL in NTF and TF cells when incubated with THZ1. For this experiment, we used a concentration of 300 nM THZ1 for 2 h (Fig. 6A). Figure 6B shows that for *ID2, lncRNA LOC730101, HEXIM1* and *EPAS1*, there was a clear increase in the mRNA levels in NTF cells incubated with THZ1 (Fig. 6B). In TF and NTF cells, the mRNA levels of *ID2* and the lncRNA increased and similar behaviour of *EPAS1* as that seen with TPL incubation was observed (Fig. 6B). In contrast, no clear increase in the *HEXIM1* transcript level was observed in TF cells, and in all cell types, *CRY2* level remained the same in THZ1-treated cells. These results suggest that there are some similarities in the gene expression responses to TPL and THZ1 in NTF and TF cells, but that the responses are not identical.

To explore how similar the gene expression responses to TPL and THZ1 are, we used public RNA-Seq data from an oesophageal cancer cell line treated with THZ1 (64). We found that 94 genes upregulated by TPL that correlated with an increase in the occupancy of RNAPII in the oncogenesis model used here were also upregulated by THZ1 in the oesophageal cancer cell line (Fig 6C). These results suggest that in response to transcriptional stress by either TPL or THZ1, a similar set of genes is overexpressed.

### Depletion of transcripts encoded by genes overexpressed in response to transcriptional stress augments the cells sensitivity to triptolide

The fact that some genes are overexpressed in response to TPL underscored the possibility that some of them were targets for tumour cell killing. To explore the likelihood that depleting the transcripts of genes overexpressed in response to TPL potentiates the killing effect of TPL, we chose to evaluate the effect of silencing *ID2, CRY2, HEXIM1, LOC730101* and *EPAS1* on cell viability and proliferation. NTF and TF cells were transfected with pooled siRNAs against each of these transcripts for 24, 48 or 72 hours, followed by incubation with 125 nM TPL for 4 h. Consistent with data shown above, 125 nM TPL alone or in combination with the scrambled siRNAs did not affect NTF and TF cells survival (Fig. 7, middle panels). All targeted RNAs were effectively silenced by the corresponding siRNAs, with the maximum reduction at 72 h post-transfection, as evaluated by immnunoblot and flow cytometry (Fig. 7A-E, left panels). Depletion of the mRNAs encoding for *ID2*, *CRY2* and *HEXIM1* significantly reduced the viability of TPL-treated NTF and TF cells as compared to cells incubated with scrambled siRNAs and TPL or TPL alone (Fig. 7A-C, middle panels). Different to *ID2, CRY2, HEXIM1* depletion, silencing lncRNA *LOC730101* resulted in diminished cell viability, even in the absence of TPL, but the effect of silencing this lncRNA, intensified the killing capacity of TPL in both NTF and TF cells (Fig. 7E, middle panel). Surprisingly, reducing the levels of *EPAS1* (*HIF-2A*) RNA did not result in enhanced killing capacity of TPL (Fig. 7D, middle panel). Unlike NTF cells, the majority of surviving TF cells underwent at least one round of proliferation less than NTF transformed cells following RNA silencing of all genes (including *EPAS1*) and TPL treatment, or even two. (Fig. 7A-E, right panels), In summary, these results show that depletion of some genes overexpressed in response to TPL sensitizes cells to TPL treatment; therefore, these genes are possible targets to enhance the killing effect of TPL on cancer cells.

**Figure 7.**
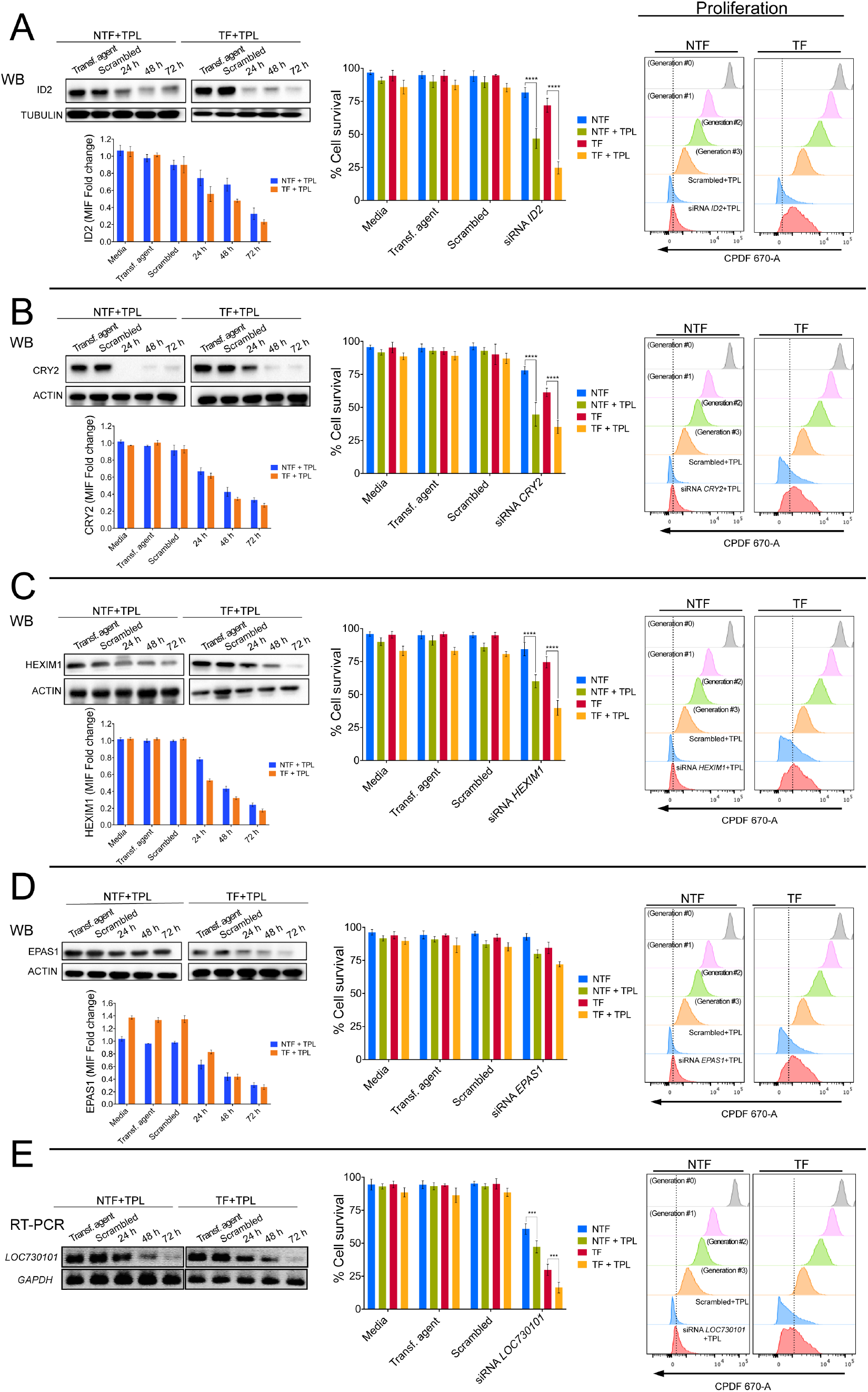
Depletion of transcripts of overexpressed genes by transcriptional stress enhances the sensitivity to TPL. *ID2, CRY2, HEXIM1 LOC73101* and *EPAS1* were depleted by incubating the corresponding siRNAs at different times. After that, the cells were treated with 125 nM of TPL for 4 h. The depletion of each protein product was determined by western blot analysis and by cytometry. In the case of the *LOC73101* transcript its depletion was analysed by RT-PCR. The effect on cell survival and proliferation were also analysed by flux cytometry and the results indicated in the figure. **(A)** ID2 analysis **(B)** CRY2 analysis **(C)** HEXIM1 analysis **(D)** EPAS analysis **(E)** LOC730101 analysis. Cell survival and proliferation only shows the effect of TPL after incubation of the corresponding siRNA by 72 h. The name of each gene is indicated in the corresponding panel.

## Discussion

The use of different chemical and physical agents is still the most common approach for killing cancer cells. The mechanism of action of chemotherapy drugs will directly kill cancer cells or stop their proliferation by inducing a cellular response linked to the mechanism of action of the drug. Substances such as TPL and THZ1 that directly impact the activities of TFIIH have a high potential for use in cancer treatment. In this study, we performed a systematic analysis of the effects and response to these drugs, primarily TPL, in the MCF10A-ErSrc oncogenesis model. Our results strongly suggest that TF tumour cells exhibited increased sensitivity to TPL or THZ1 as compared to their NTF counterparts and that the combination of both drugs had an additive effect on cell death. From a mechanistic point of view, we found that even though TPL and THZ1 directly affect transcription initiation by RNAPII of the majority of genes, specific genes are overexpressed as a result of the transcriptional stress to which the cells are submitted upon treatment with these drugs, underscoring the possibility to target these genes in conjunction with TFIIH inhibitors to kill cancer cells.

In this oncogenic model, both TPL and THZ1 were found to kill cells via apoptosis, stop proliferation one replication cycle earlier in TF cells than in NTF cells and arrest cells in G_1_. As expected, co-treatment with TPL and THZ1 had an additive effect on cell viability. Interestingly, the combination of TPL and THZ1 killed preferentially the transformed (cancerous) cells with high efficacy. These results also suggest that the simultaneous use of substances that affect different TFIIH functions is an interesting alternative to treat cancer and opens the possibility of searching for new drugs that may affect other TFIIH subunits, particularly if we consider the hepatotoxic effect of TPL and its derivatives (65).

Inhibition of transcription by TPL induces the proteasome-dependent degradation of RNAPII (66, 67). It has been proposed that TPL induces the degradation of the polymerase following phosphorylation of POLR2A, the largest subunit of RNAPII by CDK7 (66). We report here that incubation with TPL induced the degradation not only of RNAPII but also of XPB as well as of P52 and P8, but it did not affect other TFIIH subunits. THZ1, which inhibits the kinase activity of CDK7, did not have any effect on the RNAPII and TFIIH protein levels.

It is known that the interaction of XPB with P52 and P8 modulates its ATPase activity (43,45,68), and the recently solved structure of the TFIIH core shows that the N-terminal domain of XPB and the clutch domain of P52 have a similar fold and form a symmetric dimer. This interaction is not close to the ATPase domain, suggesting that regulation of ATPase activity occurs through the combined interaction of P52 and P8 with XPB (38). Accordingly, our findings show that the covalent binding of TPL to the ATPase domain of XPB destabilizes its interaction with P52, hindering the assembly of the subunits Furthermore, our computational study allowed us to propose that the dissociation/degradation of the XPB-P52-P8 submodule is mainly caused by the separation of XPB HD1-HD2 induced by the presence of TPL at the domain interface. Additionally, these results suggest that in the context of TFIIH, XPB, P52 and P8 form a submodule that is stable only when the three subunits are interacting and that the other TFIIH modules are not affected in its absence. This idea is consistent with the conformation and organization model described in the recently solved structure of the TFIIH core (38).

Intriguingly, degradation of RNAPII, as well as the XPB-P52-P8 submodule was accelerated in TF cells incubated with TPL. Thus, it is feasible the mechanism that increases the sensitivity of TF cells to TPL is partially due to the high proliferation rate of these cells, the turnover of these proteins is not fast enough to compensate for RNAPII depletion and, as a consequence, the effect on global transcription is enhanced.

Several reports on the effect of TPL using PRO-Seq and GRO-Seq have shown that the immediate effect is the reduction in transcription initiation of at least 90% of genes (21, 22). These studies were performed after short incubation times, with extremely high concentrations (10-500 µM) of TPL, but when the levels of RNAPII were not affected. We analysed the transcriptome of NTF and TF cells under conditions in which the levels of RNAPII were reduced by half, without affecting cell viability, however we detected that many transcripts were downregulated and many were upregulated. These results correlate with our data in *Drosophila,* in which both up- and downregulated transcripts were observed in P8 and P52 mutant organisms (47, 48). Interestingly, the increase in the levels of several specific genes in response to TPL also occurred in other breast cancer cell lines. In similarity with our results, it has been documented that UV irradiation causes the degradation of the RNAPII, but with the remaining RNAPII, the cell overexpress specific genes as response to this insult. A similar situation may be operating as a response to TPL (69). Furthermore, our ChIP-Seq results showed that the increase in the RNA levels of many genes was correlated with an increase in the occupancy of RNAPII on the corresponding promoters as well as in the gene bodies and qRT-PCR of intronic sequences confirmed that it is due to an enhancement of transcription. This is supported by the fact that the corresponding protein products of the genes analysed here also increase after the TPL treatment. An explanation for this phenomenon could be that, as recently reported (70), TPL inhibits transcription by its interaction with XPB, but if XPB is not present, then transcription is not inhibited. In support of this hypothesis, we found that TPL induced degradation of the XPB-P52-P8 submodule. Although it is feasible that under our experimental conditions, some genes were not affected by TPL, probably because some genes do not require XPB as it has been shown in yeast (15), this hypothesis is not supported by the observation that the response is the overexpression of specific genes, with a significant increase in the corresponding RNA levels and protein products. In addition, we observed that numerous genes upregulated by TPL were also upregulated in cells treated with THZ1, which inhibits CDK7 but does not affect the levels of the TFIIH subunits or RNAPII. Therefore, our data evidence that TPL and THZ1 activate a mechanism of gene response to transcriptional stress in treated cells.

TPL is a very effective substance for killing cancer cells, and related compounds are now in clinical phases of development. However, one of the problems with the use of TPL in patients is that it is highly toxic, and off-target effects cannot be ignored (20, 65). Therefore, finding new targets to enhance the effect of reduced concentrations of TPL is very attractive. Here, we analysed only five of many other possible targets found to be overexpressed in response to TPL and in four, its depletion enhances the toxic effect of TPL at low concentrations. These results suggest that many of the other gene products overexpressed in response to TPL may improve the anti-tumoural capacity of TPL, opening a new avenue to complement the attack on the transcriptional addiction of cancer cells.

Although treatment with TPL at high doses eventually killed most TF cells, the circumstances may be different under i*n vivo* conditions, and it is possible that many of the cancerous cells in the tumour are exposed to lower concentrations of TPL, allowing them to respond to transcriptional stress, by upregulating the transcription level of some of the genes we report. This gene response is relevant for the treatment of tumours by TPL or THZ1, more so considering that some genes that we found to be over-activated encode factors that promote tumourigenesis and/or metastasis.

In conclusion, the results presented here show that cells have the capacity to respond to the transcriptional insult caused by TPL by overexpressing some genes. Some of these genes are also overexpressed in response to THZ1, and these genes are possible targets in combination with TPL to preferentially kill cancer cells. However, this study also invokes several questions. For instance, does the depletion of genes overexpressed in response to transcriptional stress also enhance the effect of THZ1 on cancer cells? What is the mechanism that potentiates the effect of TPL by depleting genes that have different functions, such as *ID2, HEXIM1* and *CRY2*? Is the overexpression of some genes by TPL is achieved via only one specific response pathway, or are many pathways involved? The answers to these questions will be relevant in understanding the response to chemotherapy based on transcription inhibitors used in cancer.

## Aknowledgments

This work was supported by the CONACyT grant No. 1977-Problemas Nacionales program, PAPPIT/UNAM grant No. IN200315 and CONACyT No. 25088 to MZ. M.U.-A. received a scholarship from CONACyT (414127), as a student of the Programa de Doctorado en Ciencias Bioquímicas at the Universidad Nacional Autónoma de México. We thank Arturo Pimentel, Andres Saralegui, Chris Wood and the National Laboratory of Advance Microscopy at the Instituo de Biotecnología/UNAM for advice in the use of the confocal microscopes

---

**Supplementary Figure 1.**
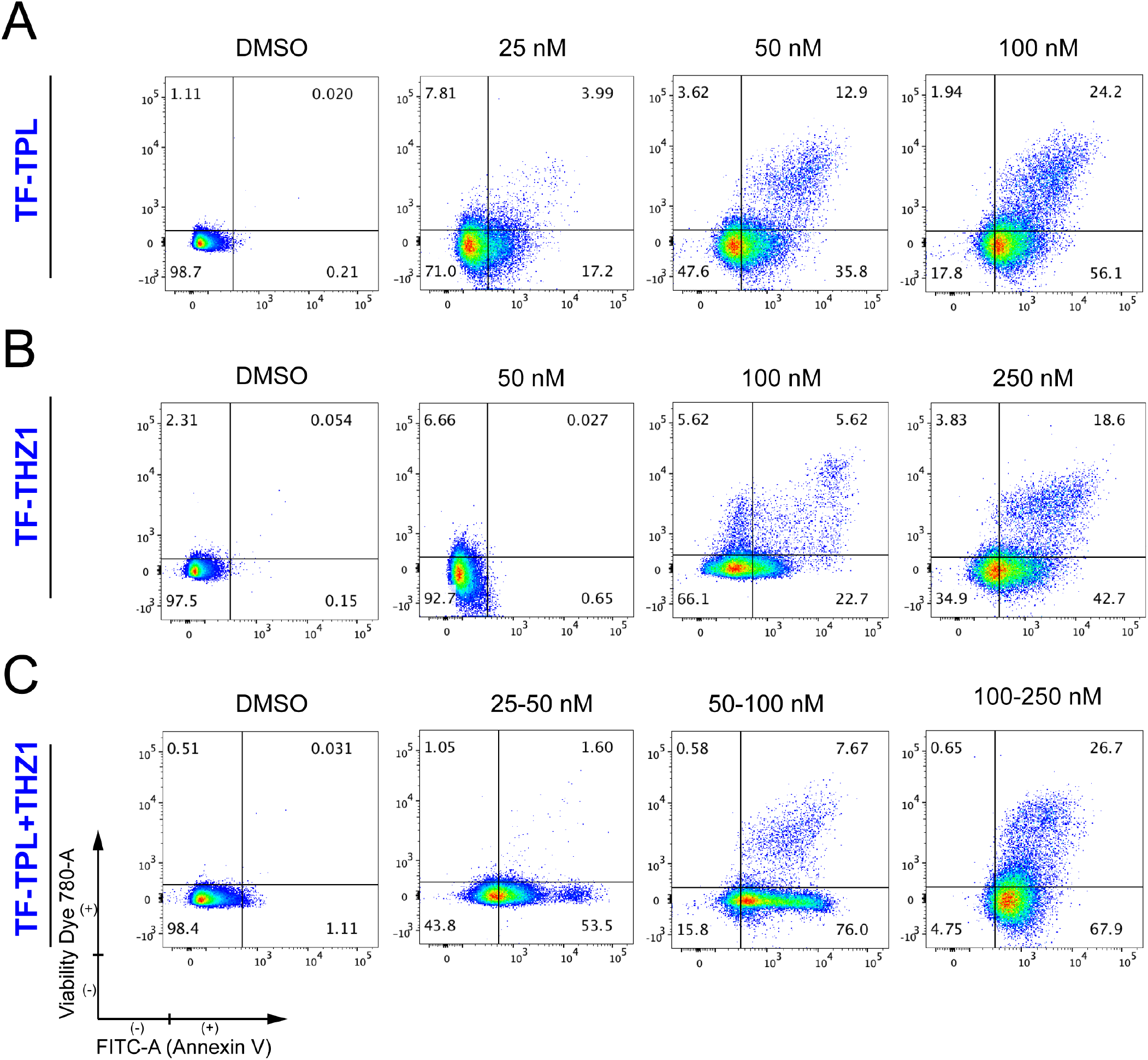
Triptolide (TPL), THZ1 and TPL/THZ1 combinatory, treatments induce apoptosis in MCF10A-ErSrc cells. Apoptosis was determined by Annexin-FITC (x-axis) and cell viability by Viability Dye 780 (y-axis). **(A)** TF cells treated with TPL for 72 h at 25, 50 and 100 nM. **(B)** TF cells treated with THZ1 for 72 h at 50, 100 and 250 nM. **(C)** TPL/THZ1 combinatory treatment, 25/50, 50/100 and 100/250 nM of each substance.

**Supplementary Figure 2.**
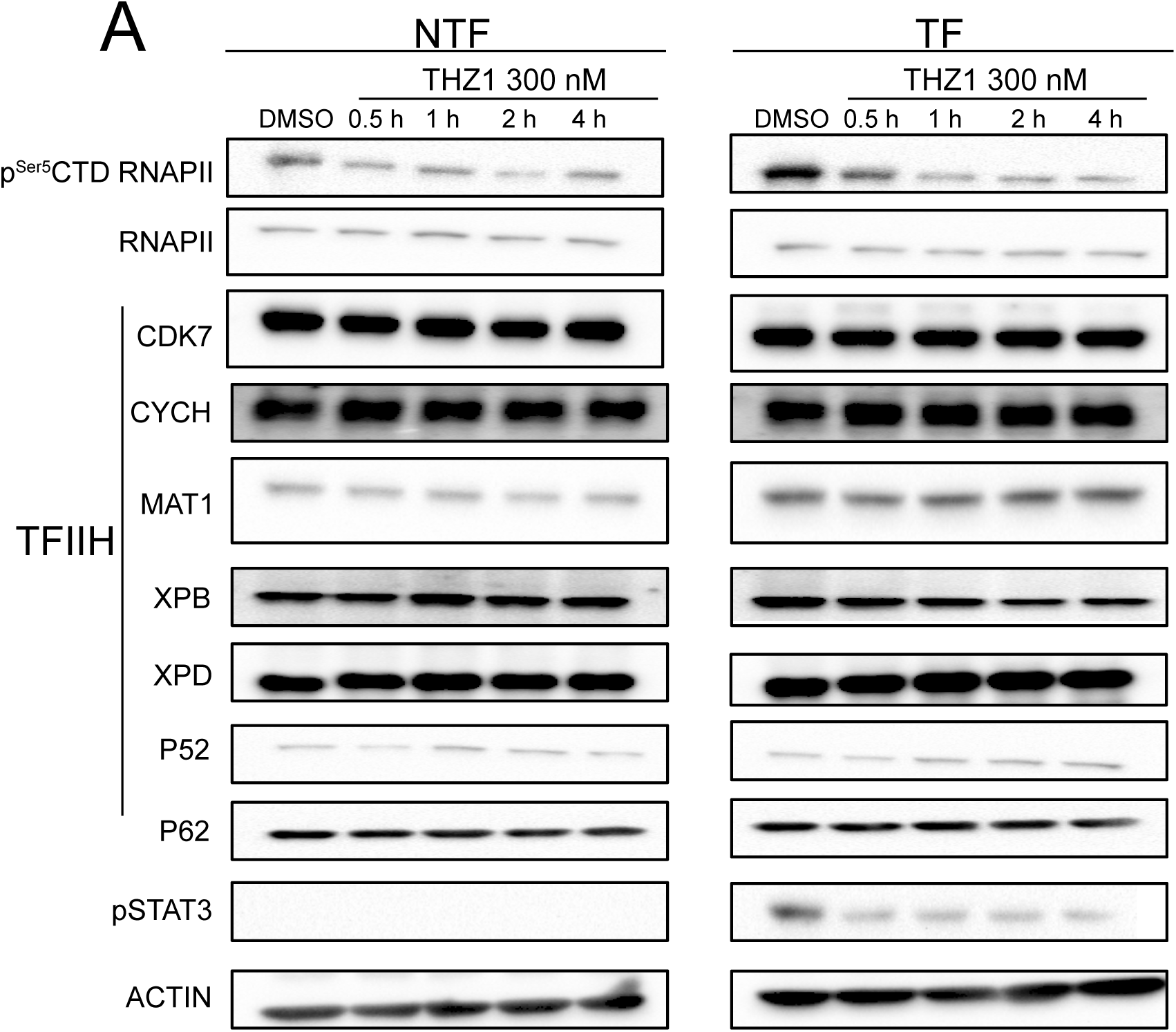
Effect of THZ1 on TFIIH. **(A)** Effect of THZ1 in NTF and TF cells treated with THZ1 with 300 nM at 0.5, 1, 2 and 4 h in comparison with the control with DMSO for 4 h. The levels of RNAPII CTD and p^Ser5^CDT RNAPII were evaluated as well as some TFIIH subunits (CDK7, CYCH, MAT1, XPB, XPD, P52, and P62). The p-STAT3 is used as control of transformation in the line and actin protein as charge control. In all the analysis the n=3.

**Supplementary Figure 3.**
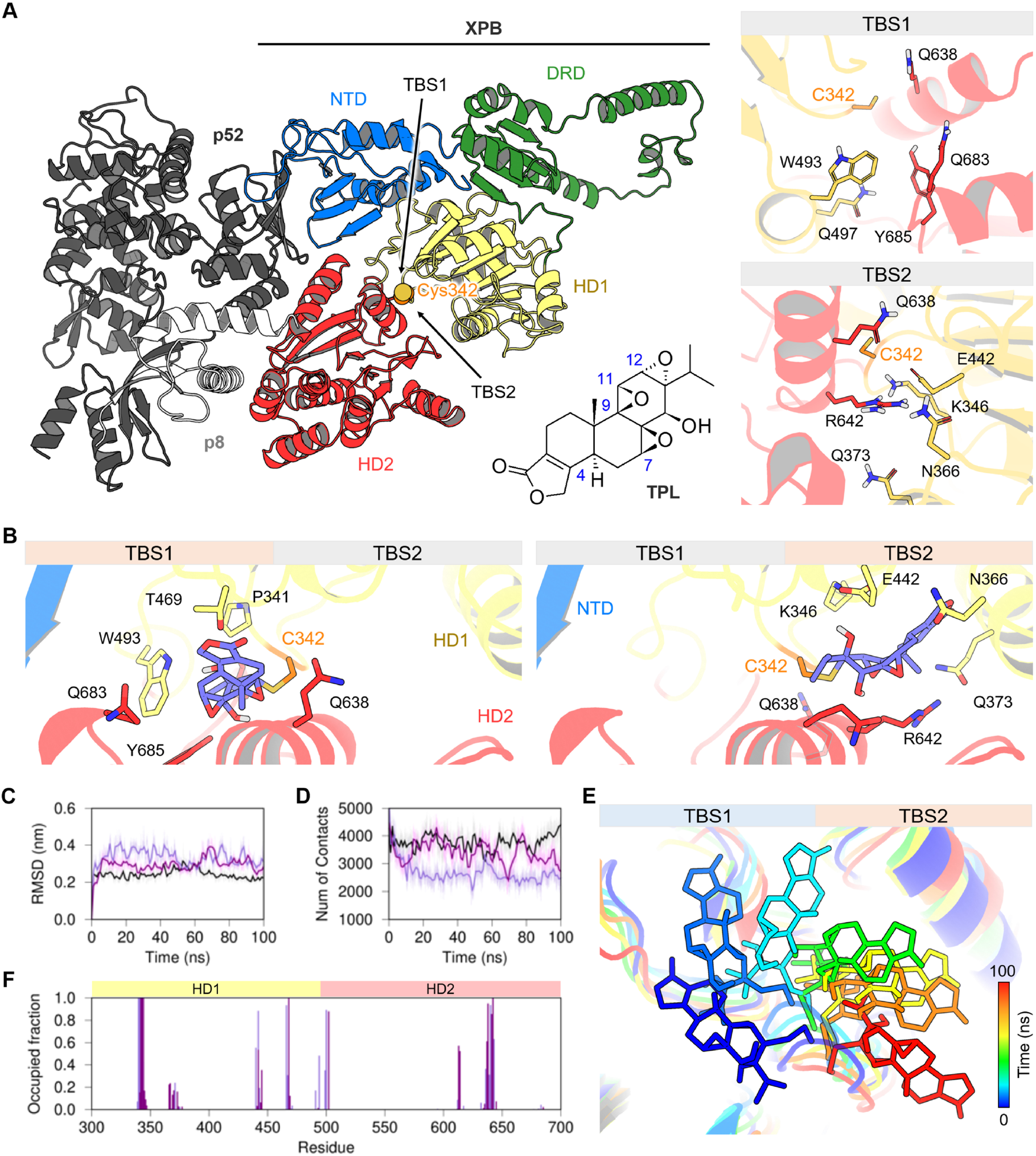
XPB-P52-P8 submodule of TFIIH and TPL interactions. **(A)** Depiction of the XPB-P52-P8 submodule tridimensional structure, the XPB component is colored by its domains: NTD (blue), DRD (green), HD1 (yellow) and HD2 (red). TPL chemical structure at the bottom shows the carbon numbering of the five potential attack sites by C342. Panel on the right show the amino acid residues that constitute the TBS1 (upper) and TBS2 (bottom) at the HD1-HD2 interface of XPB. **(B)** Best-ranked binding poses of TPL covalently bound to C342 residue at the two possible binding sites: TBS1 (left) and TBS2 (right). **(C)** The backbone root-mean-square deviations (RMSD) of apo (black) and the TBS1 (light purple) and TBS2 (dark purple) TPL-bound structures of XPB subunit. **(D)** Number of contacts formed between the HD1 and HD2 domains of XPB in the apo and the two bound states during the simulation. **(E)** Depiction of TPL shift from TBS1 to TBS2 through 100 ns MD simulation. **(F)** Occupied fraction of TPL contacts with the HD1 and HD2 domains during the last 80 ns of the simulated time.

**Supplementary Figure 4.**
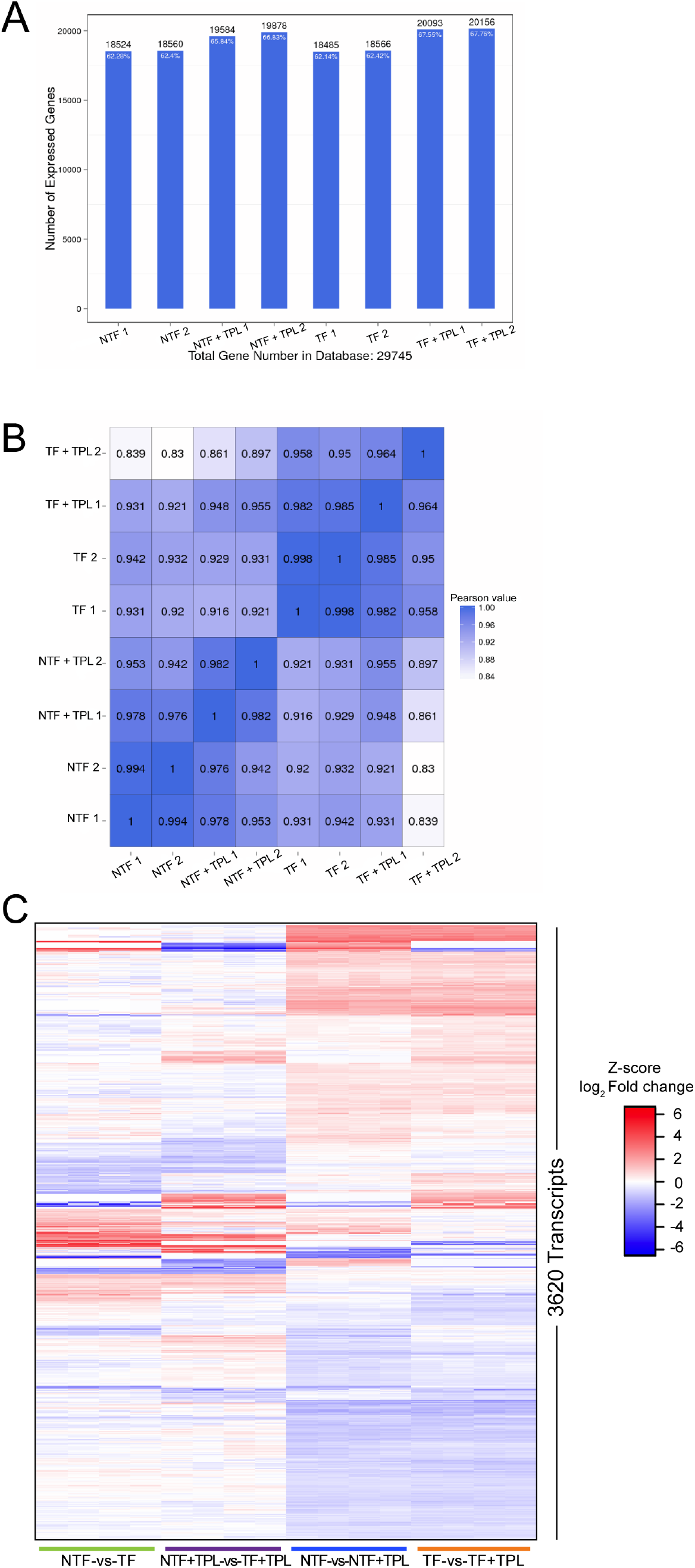
Global correlation between the expressed genes obtained in the transcriptional analysis. **(A)** Number of genes identified in the transcriptional analysis, in the x-axis is the sample name and in y-axis are the identified expressed genes. The proportion at the top of each bar represents the number of expressed genes number divided by the total gene number reported in the database. **(B)** Correlation coefficient values across each sample; the barcode colour gradient indicates the minimum value as white and the maximum as blue. If one sample is highly similar to another one, the correlation value between them is very close to 1. **(C)** Genes that were differentially expressed (3620 transcripts) in all pairwise of cluster plans.

**Supplementary Figure 5.**
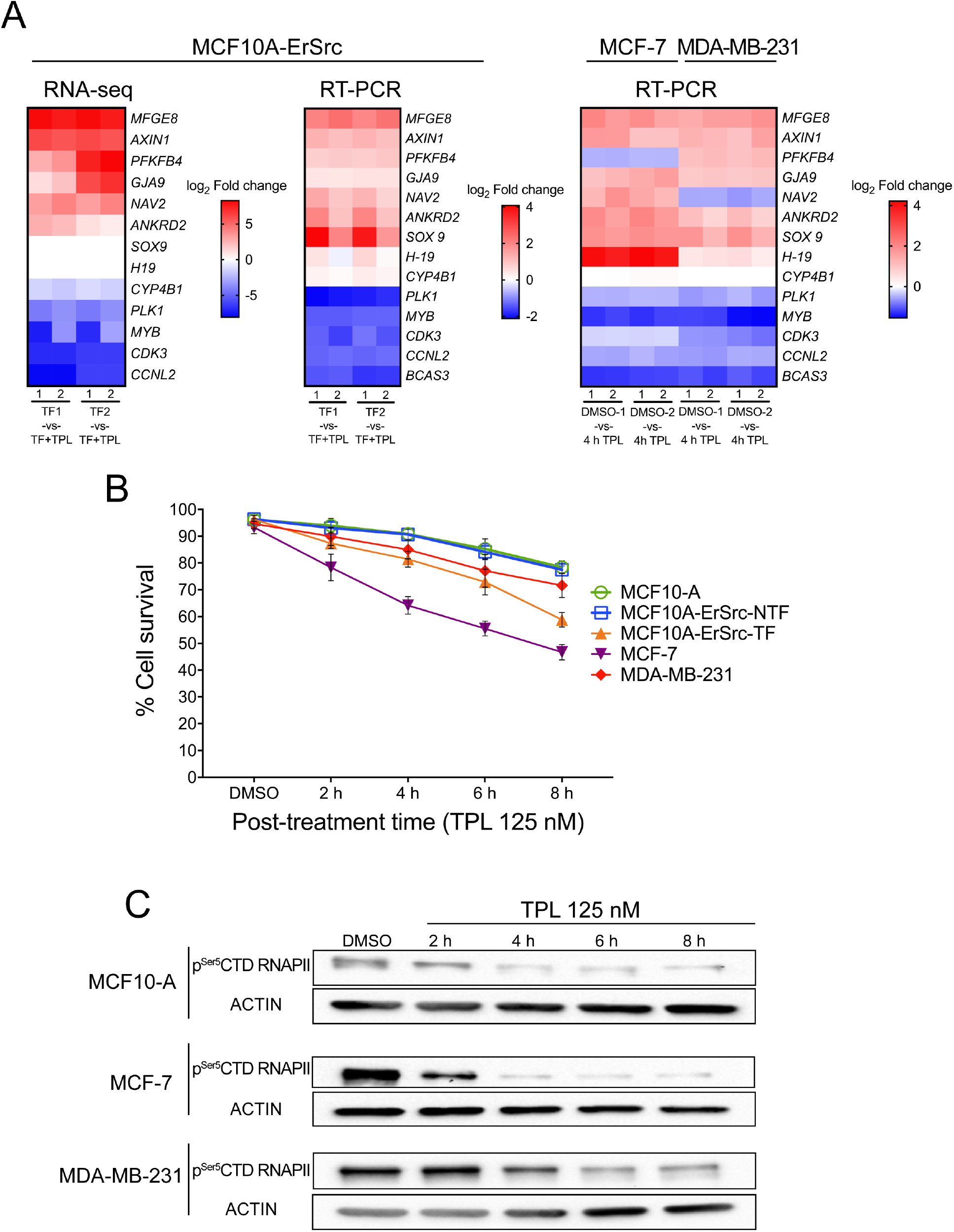
Transcriptional data of selected genes to corroborate TPL effect on transformed cells. **(A)** Left panel shows the expression levels from data obtained by the RNA-Seq. Central panel shows the expression levels of the same genes verified by RT-PCR and right panel shows the expression levels of the same genes analysed in other cell lines **(B)** Cell viability of MCF10-A (green), MCF10A-ERSrc-NTF (blue), MCF10A-ERSrc-TF (orange), MCF-7 (Purple) and MDA-MB-231 (red) cell lines are treated with TPL at 125 nM for 2, 4, 6 and 8 h. The control cells were treated with growth medium or DMSO for 8 h. **(C)** p^Ser5^CTD RNAPII levels in the MCF10-A, MCF-7, and MDA-MD-231 cell lines, after being incubated with TPL. Control cells were incubated with DMSO for 8 h.

**Supplementary Figure 6.**
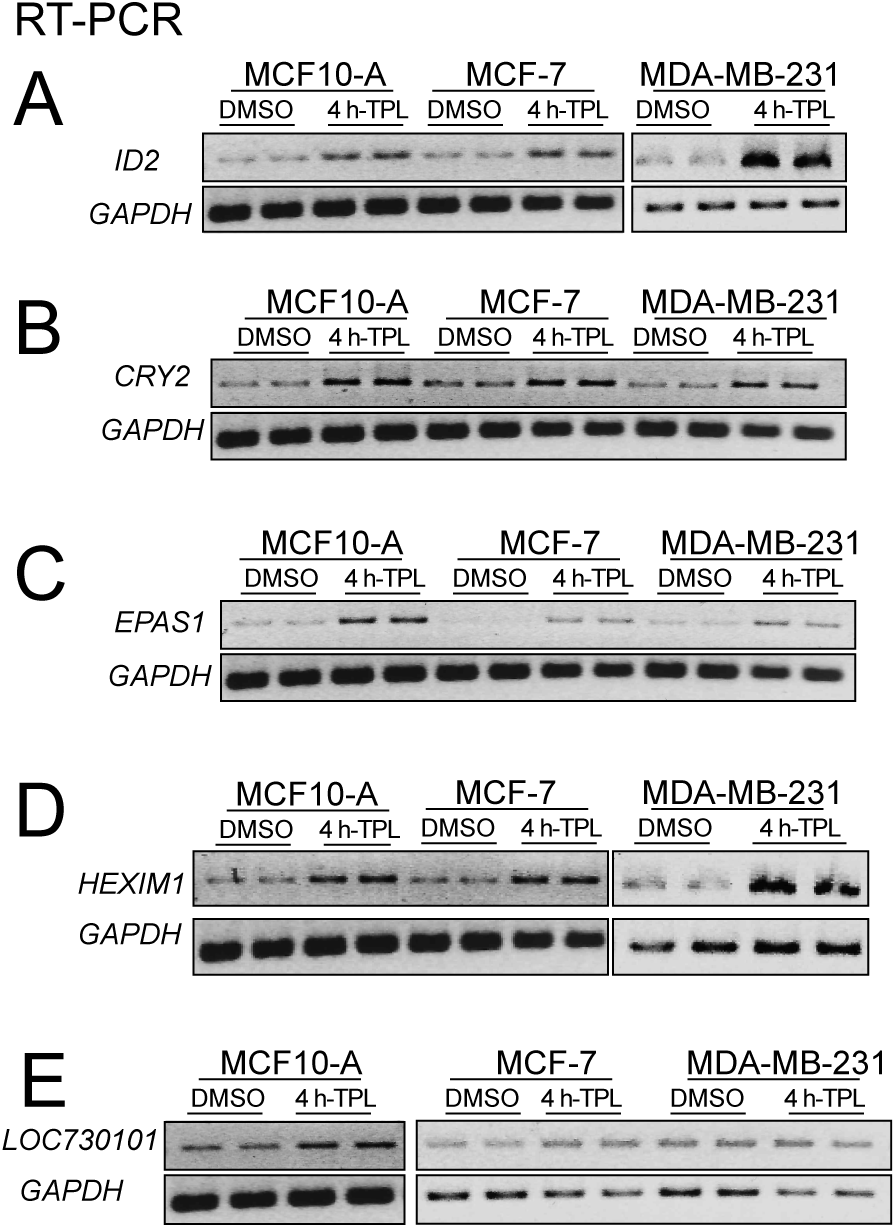
Analysis of genes overexpressed in response to TPL in other breast cancer cell lines. **(A)** ID2 RT-PCR analysis of the ID2 in the MCF10A, MCF7 and MDA-MB-231 cell lines. **(B)** LOC730101 RT-PCR analysis in the MCF10A, MCF7 and MDA-MB-231 cell lines. **(C)** HEXIM1 RT-PCR analysis in the MCF10A, MCF7 and MDA-MB-231 cell lines. **(D)** CRY2 RT-PCR analysis in the MCF10A, MCF7 and MDA-MB-231 cell lines. **(E)** EPAS1 RT-PCR analysis in the MCF10A, MCF7 and MDA-MB-231 cell lines.

## References

1. Bradner JE, Hnisz D, Young RA. Review Transcriptional Addiction in Cancer. Cell [Internet]. Elsevier Inc.; 2017;168:629–43. Available from: http://dx.doi.org/10.1016/j.cell.2016.12.013

2. Villicaña C, Cruz G, Zurita M. The basal transcription machinery as a target for cancer therapy. Cancer Cell Int. 2014;14.

3. Schilbach S, Hantsche M, Tegunov D, Dienemann C, Wigge C, Urlaub H, et al. Structures of transcription pre-initiation complex with TFIIH and Mediator. Nature. 2017;

4. Adelman K, Lis JT. Promoter-proximal pausing of RNA polymerase II: emerging roles in metazoans. Nat Rev Genet [Internet]. Nature Publishing Group; 2012;13:720–31. Available from: http://dx.doi.org/10.1038/nrg3293

5. Liu X, Kraus WL, Bai X. Ready, pause, go: Regulation of RNA polymerase II pausing and release by cellular signaling pathways. Trends Biochem Sci [Internet]. Elsevier Ltd; 2015;40:516–25. Available from: http://dx.doi.org/10.1016/j.tibs.2015.07.003

6. Bowman EA, Kelly WG. RNA Polymerase II transcription elongation and Pol II CTD Ser2 phosphorylation: A tail of two kinases. Nucl (United States). 2014;5.

7. Jonkers I, Lis JT. Getting up to speed with transcription elongation by RNA polymerase II. Nat Rev Mol Cell Biol [Internet]. Nature Publishing Group; 2015;16:167–77. Available from: http://dx.doi.org/10.1038/nrm3953

8. Berico P, Coin F. Is TFIIH the new Achilles heel of cancer cells? Transcription [Internet]. Taylor & Francis; 2018;9:47–51. Available from: https://doi.org/10.1080/21541264.2017.1331723

9. Zurita M, Cruz-Becerra G. TFIIH: New discoveries regarding its mechanisms and impact on cancer treatment. J Cancer. 2016;7:2258–65.

10. Compe E, Egly J-M. Nucleotide Excision Repair and Transcriptional Regulation: TFIIH and Beyond. Annu Rev Biochem. 2016;85:265–90.

11. Compe E, Egly JM. TFIIH: When transcription met DNA repair. Nat Rev Mol Cell Biol. 2012;13:343–54.

12. Fisher RP. Cdk7: a kinase at the core of transcription and in the crosshairs of cancer drug discovery. Transcription [Internet]. Taylor & Francis; 2019;10:47– 56. Available from: https://doi.org/10.1080/21541264.2018.1553483

13. Coin F, Egly JM. Revisiting the Function of CDK7 in Transcription by Virtue of a Recently Described TFIIH Kinase Inhibitor. Mol. Cell. 2015.

14. Nilson KA, Guo J, Turek ME, Brogie JE, Delaney E, Luse DS, et al. THZ1 Reveals Roles for Cdk7 in Co-transcriptional Capping and Pausing. Mol Cell [Internet]. Elsevier Inc.; 2015;59:576–87. Available from: http://dx.doi.org/10.1016/j.molcel.2015.06.032

15. Dienemann C, Schwalb B, Schilbach S, Cramer P. Promoter Distortion and Opening in the RNA Polymerase II Cleft. Mol Cell. 2019;73:97–106.e4.

16. Greber BJ, Nguyen THD, Fang J, Afonine P V., Adams PD, Nogales E. The cryo-electron microscopy structure of human transcription factor IIH. Nature. 2017;

17. Christensen CL, Kwiatkowski N, Abraham BJ, Carretero J, Al-Shahrour F, Zhang T, et al. Targeting Transcriptional Addictions in Small Cell Lung Cancer with a Covalent CDK7 Inhibitor. Cancer Cell. 2014;26:909–22.

18. Zeng M, Kwiatkowski NP, Zhang T, Nabet B, Xu M, Liang Y, et al. Targeting MYC dependency in ovarian cancer through inhibition of CDK7 and CDK12/13. Elife. 2018;7:1–20.

19. Titov D V., Gilman B, He QL, Bhat S, Low WK, Dang Y, et al. XPB, a subunit of TFIIH, is a target of the natural product triptolide. Nat Chem Biol [Internet]. Nature Publishing Group; 2011;7:182–8. Available from: http://dx.doi.org/10.1038/nchembio.522

20. Noel P, Von Hoff DD, Saluja AK, Velagapudi M, Borazanci E, Han H. Triptolide and Its Derivatives as Cancer Therapies. Trends Pharmacol Sci [Internet]. Elsevier Ltd; 2019;40:327–41. Available from: https://doi.org/10.1016/j.tips.2019.03.002

21. Henriques T, Gilchrist DA, Nechaev S, Bern M, Muse GW, Burkholder A, et al. Stable pausing by RNA polymerase II provides an opportunity to target and integrate regulatory signals. Mol Cell [Internet]. Elsevier Inc.; 2013;52:517–28. Available from: http://www.ncbi.nlm.nih.gov/pubmed/24184211 http://www.pubmedcentral.nih.gov/articlerender.fcgi?artid=PMC3845087

22. Jonkers I, Kwak H, Lis JT. Genome-wide dynamics of Pol II elongation and its interplay with promoter proximal pausing, chromatin, and exons. Elife. 2014;3:1– 25.

23. Rimel JK, Taatjes DJ. The essential and multifunctional TFIIH complex. 2018;27:1018–37.

24. Iliopoulos D, Hirsch HA, Struhl K. An Epigenetic Switch Involving NF-κB, Lin28, Let-7 MicroRNA, and IL6 Links Inflammation to Cell Transformation. Cell [Internet]. Elsevier Ltd; 2009;139:693–706. Available from: http://dx.doi.org/10.1016/j.cell.2009.10.014

25. Debnath J, Muthuswamy SK, Brugge JS. Morphogenesis and oncogenesis of MCF-10A mammary epithelial acini grown in three-dimensional basement membrane cultures. Methods. 2003;30:256–68.

26. Neve RM, Chin K, Fridlyand J, Yeh J, Baehner FL, Fevr T, et al. A collection of breast cancer cell lines for the study of functionally distinct cancer subtypes. Cancer Cell. 2006;10:515–27.

27. Cabantous S, Nguyen HB, Pedelacq JD, Koraïchi F, Chaudhary A, Ganguly K, et al. A new protein-protein interaction sensor based on tripartite split-GFP association. Sci Rep. 2013;3:1–9.

28. Gurrion C, Uriostegui M, Zurita M. Heterochromatin reduction correlates with the increase of the KDM4B and KDM6A demethylases and the expression of pericentromeric DNA during the acquisition of a transformed phenotype. J Cancer. 2017;8:2866–75.

29. Schmittgen TD, Livak KJ. Analyzing real-time PCR data by the comparative C T method. 2008;3:1101–8.

30. Holmes KL, Otten G, Yokoyama WM. Flow Cytometry Analysis Using the Becton Dickinson FACS Calibur. 2001;1–22.

31. Langmead B, Trapnell C, Pop M, Salzberg SL. Ultrafast and memory-efficient alignment of short DNA sequences to the human genome. 2009;10.

32. Li H, Durbin R. Fast and accurate short read alignment with Burrows – Wheeler transform. 2009;25:1754–60.

33. Li B, Dewey CN. RSEMD: accurate transcript quantification from RNA-Seq data with or without a reference genome. 2011;

34. Liang K, Volk AG, Haug JS, Marshall SA, Woodfin AR, Bartom ET, et al. Therapeutic Targeting of MLL Degradation Pathways in MLL-Rearranged Leukemia. Cell [Internet]. Elsevier; 2017;168:59–72.e13. Available from: http://dx.doi.org/10.1016/j.cell.2016.12.011

35. Li R, Yu C, Li Y, Lam T, Yiu S, Kristiansen K, et al. SOAP2D: an improved ultrafast tool for short read alignment. 2009;25:1966–7.

36. Zhang Y, Liu T, Meyer CA, Eeckhoute J, Johnson DS, Bernstein BE, et al. Open Access Model-based Analysis of ChIP-Seq (MACS). 2008;

37. Heinz S, Benner C, Spann N, Bertolino E, Lin YC, Laslo P, et al. Article Simple Combinations of Lineage-Determining Transcription Factors Prime cis - Regulatory Elements Required for Macrophage and B Cell Identities. MOLCEL [Internet]. Elsevier; 2010;38:576–89. Available from: http://dx.doi.org/10.1016/j.molcel.2010.05.004

38. Greber BJ, Toso DB, Fang J, Nogales E. The complete structure of the human TFIIH core complex. Elife. 2019;8:1–29.

39. Bianco G, Forli S, Goodsell DS, Olson AJ. Covalent docking using autodock: Two-point attractor and flexible side chain methods. Protein Sci. 2016;25:295– 301.

40. Lindorff-Larsen K, Piana S, Palmo K, Maragakis P, Klepeis JL, Dror RO, et al. Improved side-chain torsion potentials for the Amber ff99SB protein force field. Proteins Struct Funct Bioinforma. 2010;78:1950–8.

41. Abraham MJ, Murtola T, Schulz R, Páall S, Smith JC, Hess B, et al. GROMACS: High performance molecular simulations through multi-level parallelism from laptops to supercomputers. SoftwareX. 2015;1–2:19–25.

42. Iliopoulos D, Rotem A, Struhl K. Inhibition of miR-193a expression by max and RXRα activates K-ras and PLAU to mediate distinct aspects of cellular transformation. Cancer Res. 2011;71:5144–53.

43. Fregoso M, Laine J-P, Aguilar-Fuentes J, Mocquet V, Reynaud E, Coin F, et al. DNA Repair and Transcriptional Deficiencies Caused by Mutations in the Drosophila p52 Subunit of TFIIH Generate Developmental Defects and Chromosome Fragility. Mol Cell Biol. 2007;27:3640–50.

44. Kwiatkowski N, Zhang T, Rahl PB, Abraham BJ, Reddy J, Ficarro SB, et al. Targeting transcription regulation in cancer with a covalent CDK7 inhibitor. Nature [Internet]. Nature Publishing Group; 2014;511:616–20. Available from: http://dx.doi.org/10.1038/nature13393

45. Coin F, Oksenych V, Egly JM. Distinct Roles for the XPB/p52 and XPD/p44 Subcomplexes of TFIIH in Damaged DNA Opening during Nucleotide Excision Repair. Mol Cell. 2007;26:245–56.

46. He QL, Titov D V., Li J, Tan M, Ye Z, Zhao Y, et al. Covalent modification of a cysteine residue in the XPB subunit of the general transcription factor TFIIH through single epoxide cleavage of the transcription inhibitor triptolide. Angew Chemie - Int Ed. 2015;

47. Cruz-Becerra G, Juárez M, Valadez-Graham V, Zurita M. Analysis of Drosophila p8 and p52 mutants reveals distinct roles for the maintenance of TFIIH stability and male germ cell differentiation. Open Biol. 2016;6.

48. Cruz-Becerra G, Valerio-Cabrera S, Juárez M, Bucio-Mendez A, Zurita M. TFIIH is highly dynamic during zygotic genome activation in Drosophila and its depletion causes catastrophic mitosis. J Cell Sci. 2018;jcs.211631.

49. Ji Z, He L, Rotem A, Janzer A, Cheng CS, Regev A, et al. Genome-scale identification of transcription factors that mediate an inflammatory network during breast cellular transformation. Nat Commun [Internet]. Springer US; 2018;9. Available from: http://dx.doi.org/10.1038/s41467-018-04406-2

50. Chen F, Gao X, Shilatifard A, Shilatifard A. Stably paused genes revealed through inhibition of transcription initiation by the TFIIH inhibitor triptolide. Genes Dev. 2015;29:39–47.

51. Erickson B, Sheridan RM, Cortazar M, Bentley DL. Dynamic turnover of paused pol II complexes at human promoters. Genes Dev. 2018;32:1215–25.

52. Hurst DR, Xie Y, Vaidya KS, Mehta A, Moore BP, Accavitti-Loper MA, et al. Alterations of BRMS1-ARID4A interaction modify gene expression but still suppress metastasis in human breast cancer cells. J Biol Chem. 2008;283:7438–44.

53. Solimini NL, Liang AC, Xu C, Pavlova NN, Xu Q, Davoli T, et al. STOP gene Phactr4 is a tumor suppressor. Proc Natl Acad Sci. 2013;110:E407–14.

54. Jiang B, Chen W, Qin H, Diao W, Li B, Cao W, et al. TOX3 inhibits cancer cell migration and invasion via transcriptional regulation of SNAI1 and SNAI2 in clear cell renal cell carcinoma. Cancer Lett [Internet]. Elsevier; 2019;449:76–86. Available from: http://www.ncbi.nlm.nih.gov/pubmed/30772441

55. Ju W, Yoo BC, Kim I-J, Kim JW, Kim SC, Lee HP. Identification of Genes With Differential Expression in Chemoresistant Epithelial Ovarian Cancer Using High-Density Oligonucleotide Microarrays. Oncol Res Featur Preclin Clin Cancer Ther. 2009;

56. Jung HY, Fattet L, Tsai JH, Kajimoto T, Chang Q, Newton AC, et al. Apical– basal polarity inhibits epithelial–mesenchymal transition and tumour metastasis by PAR-complex-mediated SNAI1 degradation. Nat Cell Biol [Internet]. Springer US; 2019;21:359–71. Available from: http://dx.doi.org/10.1038/s41556-019-0291-8

57. Lee H, Goodarzi H, Tavazoie SF, Alarcón CR. TMEM2 Is a SOX4-regulated gene that mediates metastatic migration and invasion in breast cancer. Cancer Res. 2016;76:4994–5005.

58. Matsuo T, Dat LT, Komatsu M, Yoshimaru T, Daizumoto K, Sone S, et al. Early growth response 4 is involved in cell proliferation of Small cell lung cancer through transcriptional activation of its downstream genes. PLoS One. 2014;9.

59. Zeng S, Zhang Y, Ma J, Deng G, Qu Y, Guo C, et al. BMP4 promotes metastasis of hepatocellular carcinoma by an induction of epithelial– mesenchymal transition via upregulating ID2. Cancer Lett [Internet]. Elsevier Ltd; 2017;390:67–76. Available from: http://dx.doi.org/10.1016/j.canlet.2016.12.042

60. Huber AL, Papp SJ, Chan AB, Henriksson E, Jordan SD, Kriebs A, et al. CRY2 and FBXL3 Cooperatively Degrade c-MYC. Mol Cell [Internet]. Elsevier Inc.; 2016;64:774–89. Available from: http://dx.doi.org/10.1016/j.molcel.2016.10.012

61. Li S, Hu K, Chen S, Liu S, Wang Y. High expression of long non-coding RNA LOC730101 correlates with distant metastasis and exhibits a poor prognosis in patients with osteosarcoma. 2018;4115–20.

62. Bowry A, Piberger AL, Rojas P, Saponaro M, Bowry A, Piberger AL, et al. Report BET Inhibition Induces HEXIM1- and RAD51-Dependent Conflicts between Transcription and Report BET Inhibition Induces HEXIM1- and RAD51-Dependent Conflicts between Transcription and Replication. CellReports [Internet]. ElsevierCompany.; 2018;25:2061–2069.e4. Available from: https://doi.org/10.1016/j.celrep.2018.10.079

63. Ketchart W, Ogba N, Kresak A, Albert JM, Pink JJ, Montano MM. HEXIM1 is a critical determinant of the response to tamoxifen. Oncogene [Internet]. Nature Publishing Group; 2011;30:3563–9. Available from: http://dx.doi.org/10.1038/onc.2011.76

64. Jiang YY, Lin DC, Mayakonda A, Hazawa M, Ding LW, Chien WW, et al. Targeting super-enhancer-Associated oncogenes in oesophageal squamous cell carcinoma. Gut. 2017;66:1358–68.

65. Song W, Liu M, Wu J, Zhai H, Chen Y, Peng Z. Preclinical Pharmacokinetics of Triptolide: A Potential Antitumor Drug. Curr Drug Metab. 2018;

66. Manzo SG, Zhou ZL, Wang YQ, Marinello J, He JX, Li YC, et al. Natural product triptolide mediates cancer cell death by triggering CDK7-dependent degradation of RNA polymerase II. Cancer Res. 2012;72:5363–73.

67. Wang Y, Lu J jian, He L, Yu Q. Triptolide (TPL) inhibits global transcription by inducing proteasome-dependent degradation of RNA polymerase II (Pol II). PLoS One. 2011;6.

68. Kainov DE, Vitorino M, Cavarelli J, Poterszman A, Egly JM. Structural basis for group A trichothiodystrophy. Nat Struct Mol Biol. 2008;15:980–4.

69. Williamson L, Saponaro M, Boeing S, East P, Mitter R, Kantidakis T, et al. UV Irradiation Induces a Non-coding RNA that Functionally Opposes the Protein Encoded by the Same Gene. Cell [Internet]. Elsevier; 2017;168:843–855.e13. Available from: http://dx.doi.org/10.1016/j.cell.2017.01.019

70. Alekseev S, Nagy Z, Sandoz J, Weiss A, Egly JM, Le May N, et al. Transcription without XPB Establishes a Unified Helicase-Independent Mechanism of Promoter Opening in Eukaryotic Gene Expression. Mol Cell. 2017;65:504–514.e4.

